# Census and genetic analysis of the United States marmoset population

**DOI:** 10.64898/2026.05.19.726287

**Authors:** Murillo F. Rodrigues, Philberta Leung, Jamie A. Ivy, Alexandra Stendahl, Karina Ray, Jenna Castro, Samuel M. Peterson, Ricardo C. H. del Rosario, Simon Ploesch, Joanna Malukiewicz, Katinka A. Vigh-Conrad, Benjamin N. Bimber, Marmoset Genetics Working Group, Jeffrey D. Wall, Donald F. Conrad

**Affiliations:** Division of Genetics, Oregon National Primate Research Center, OHSU, Beaverton OR, US; Independent Consultant, CO, US; Stanley Center for Psychiatric Research, Broad Institute of Harvard and MIT, Boston MA, US; Department of Biology, University of Hamburg, Germany; German Primate Center, Leibniz Institute for Primate Research, Niedersachsen, Germany; Institute of Tropical Medicine, University of São Paulo, Brazil

## Abstract

The common marmoset (Callithrix jacchus), a small monkey native to Brazil, has been used as a biomedical model in the United States (US) since the 1950s, yet the origins, genomic diversity, and population structure of current colonies remain poorly defined. Through the NIH Marmoset Coordinating Center, we registered and sampled most US research marmosets (∼2,300 living animals) and assembled pedigrees and historical records for >10,000 individuals. We present a resource of >800 whole-genome sequences, largely from US colonies. These data reveal an unexpected population structure that predates the establishment of research colonies. Indeed, this population structure mirrors variation found in marmosets across Brazil. Leveraging sequenced families, we generate the first pedigree-based recombination map and improved estimates of de novo mutation processes for this species. Our insights into genetic diversity, structure, and inbreeding will guide colony management, inform disease modelling and strengthen the marmoset’s standing as a biomedical model. Further, this work demonstrates how coordinated efforts across colonies can enable a self-sustaining “living laboratory”, supporting data sharing and well-powered studies beyond the reach of single institutions.

## Main Text

Understanding the genomes of primates is critical to studies of primate evolution and human health. Among nonhuman primates, the common marmoset (*Callithrix jacchus*) is an emerging model for biomedical research because of its small size, high reproductive output, and short lifespan (*1–4*). Notably, the marmoset model has been instrumental in elucidating the neural basis of behavior and sensory processing (*5–10*), age-related cognitive decline (*11*), neurodegenerative disorders (*12–14*), and embryonic development (*15*). Despite these advances, there is minimal understanding of genomic variation across research populations, which limits their potential as a biomedical model (*16*).

The common marmoset, a South American monkey member of the Callitrichidae family, is of evolutionary interest given its unique genetic and morphological traits (*17*). Remarkably, marmosets typically reproduce by multiple gestation, and littermates routinely exchange hematopoietic stem cells through shared placental circulation, resulting in lifelong blood chimerism, a phenomenon that is nearly unique among mammals (*18–20*). Chimerism complicates studies of genetic variation, as distinguishing variants between each twin is difficult (*21*). Marmosets also exhibit reduced body size (weighing around 300g) due to dietary and reproductive adaptations (*22*, *23*), a trait that has facilitated its adoption as a laboratory model.

Marmosets have experienced varied evolutionary pressures due to their broad ecological range (*24*). Native to northeastern Brazil, marmosets inhabit environments ranging from the humid Atlantic Forest to the Brazilian semi-arid Cerrado and Caatinga biomes (*25*). It is unclear, however, what allows the species to inhabit such distinct environments and whether a strong population structure exists in the wild (*26*). These ecological differences, and the genetic adaptations associated with them, may affect captive colony management, for instance by influencing genetic disease predispositions or dietary requirements. In the past few decades, *C. jacchus* was introduced to southeast Brazil, where it now overlaps with closely related species (*27*), including *C. penicillata*, *C. aurita* and *C. geoffroyi*. This created the opportunity for anthropogenic hybridization (*28*), potentially complicating the conservation of related species in the wild and confounding biomedical research (*29*).

Given its potential as a biomedical model, marmoset colonies and breeding centers have been created across the world in the last six decades. The first successful colony in the US was probably the one established at Tulane University in the 1960s, where marmosets were used in numerous studies ranging from endocrinology to oral health (*30*, *31*). From the late 1960s through the 1980s, Brazil strengthened legal measures aimed at protecting its native wildlife, effectively closing the legal routes for exporting marmosets (*32*). During the 1970s and 1980s, when pet trade was booming, hundreds of marmosets found their way to South Africa, where they were bred and exported throughout the world (including to research colonies) (*33*). Since then, some colonies have been maintained without the introduction of new animals, while others have exchanged captive animals to mitigate inbreeding. This complex and opaque history, marking the beginning of captive marmoset colonies for public health research, underscores the need to better understand their wild origins and the effects of captivity on marmoset genetic variation and health.

Despite extensive use of marmosets in biomedical research and their unique adaptations, genomic resources for the species remain scarce (*34–37*). Here, we address this gap by generating and compiling whole-genome sequencing data for over 800 captive marmosets. This resource was produced through a collaborative effort between animal caregivers and scientists as part of the Marmoset Coordinating Center, an initiative established to facilitate information sharing and coordinate breeding among US institutions collectively housing more than 2,500 animals. Our main goal was to characterize genomic variation across colonies to improve the utility of the marmoset model, inform colony management, and learn about the biology and evolutionary history of this unique and charismatic New World monkey.

### A genomic and pedigree resource for marmoset research

The Marmoset Coordinating Center (MCC) assembled genealogical, demographic and medical data on a total of 9,979 marmosets, of which 2,343 are currently living and housed in one of 31 North American research centers (**Table S1, Fig. 1A**).

**Figure 1.**
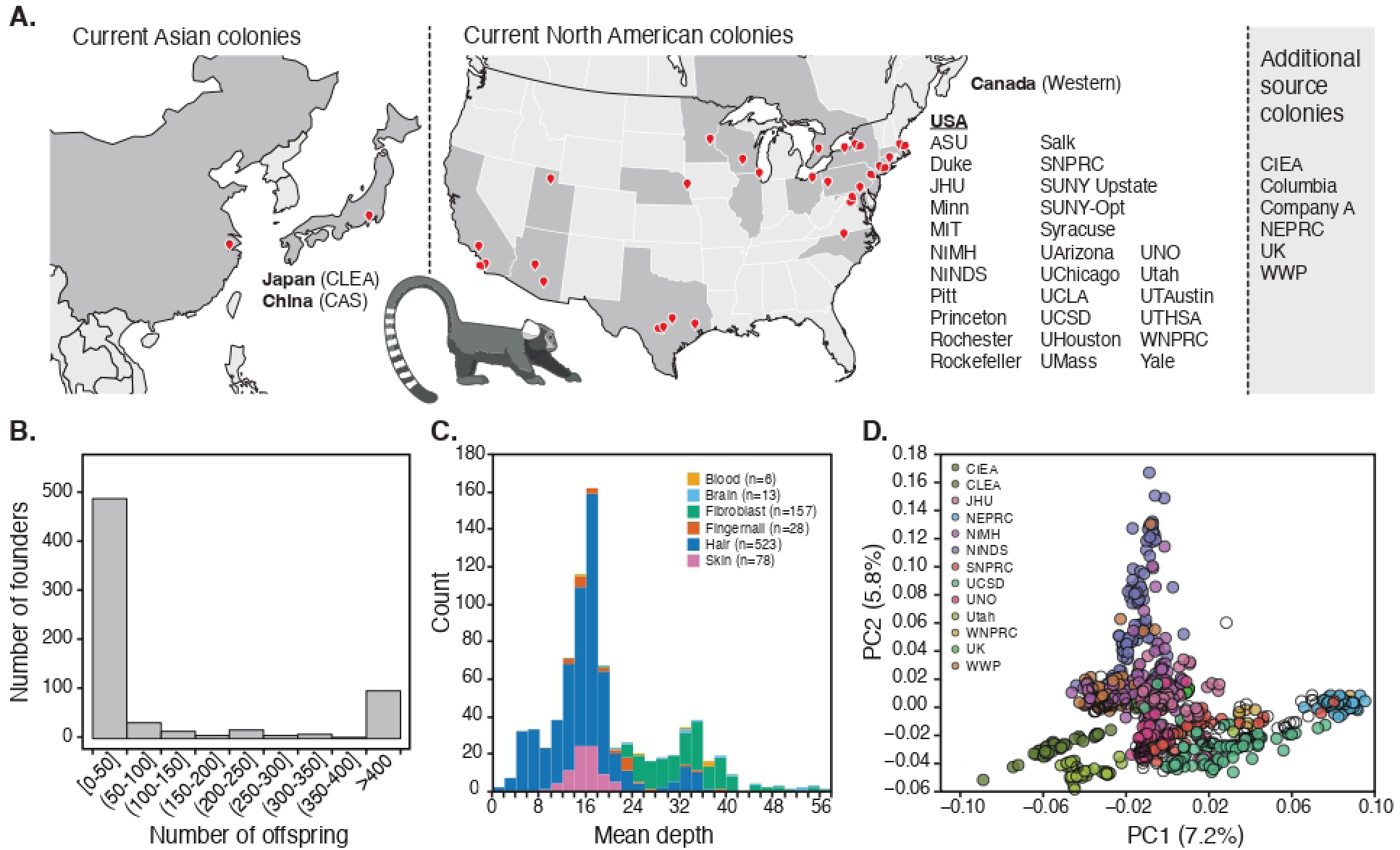
Overview of captive marmoset colonies. (A) Map with current North American and Asian colonies included in this study. Note that some animals might have been born in a di<erent colony, some of which did not contribute to the MCC or no longer exist. (B) Distribution of number of living descendants per pedigree terminal. (C) Distribution of mean sequencing depth across 805 marmoset samples analyzed in this study. (D) PCA plot based on whole-genome data for the 805 animals included in the study. Population labels are based on colony where individuals were born (source colony), where white dots include colonies with few sequenced individuals.

To our knowledge, this represents an essentially complete census of all marmosets currently being used for academic research in the US. For each of the living animals registered with the MCC, we collected 16 fields of information to construct a minimal medical and demographic profile that serves as a basis for research and husbandry. We compiled pedigree information for all animals in the MCC database, integrating all available digital and written records using the best practices adopted by zoos for studbook data entry and quality control.

Using the pedigree records, all living animals in the MCC could be traced back to 552 pedigree terminals (*i.e.*, animals with no recorded ancestors), 52% of which originated outside of the US. The average depth of the pedigree was 4.3 generations, and the average mean kinship, assuming terminal pedigree animals are unrelated, was 0.007. For context, the pedigree-based kinship of second cousins is 0.03125, suggesting that, based on the available pedigree data, captive marmosets exhibit low levels of inbreeding and have maintained relatively high genetic diversity despite the absence of a nationally coordinated breeding program. We validated the pedigree using genetic estimates of kinship, which showed good concordance with the pedigree-derived kinship coefficients, particularly for animals with good historical records (**Fig. S1**). The number of living descendants per terminal ranged from 0 to 1,132, with a median of 6 descendants per terminal (**Fig. 1B**). Across MCC-tracked animals, the sex-averaged generation time was 4.79 years (a value that will be used in downstream analyses); as in humans, the mean generation time for males was higher than females (5.11 years vs. 4.47 years) (*38*).

We also performed a genomic survey of captive marmosets, which has historically proven difficult due to the hematopoietic chimerism present in all marmoset species (*39*, *40*). To address this issue, we developed a protocol for extracting DNA from hair follicle cells (*41*). Hair follicle sampling is minimally invasive, and follicular cells do not exhibit chimerism, though small amounts of blood carried on hair follicle samples contribute to some residual chimerism, with 2.3% of reads per animal originating from twin genomes. Similarly, skin punches and fibroblast cell cultures are mostly free of chimerism (*21*), but these tissues are costlier to obtain and demand more invasive collection procedures.

We sequenced follicle DNA from 578 animals to a target of 30X depth and jointly processed the resulting data with whole genome sequencing (WGS) data from another 193 MCC animals derived from cultured fibroblasts and skin punches, as well as 34 genomes from captive marmosets from Asia (*35*, *36*) (**Fig. 1C**). We also included data from an additional *C. jacchus* trio, and two outgroup species, *C. geoffroyi* and *C. kuhli* (*42*). In some analyses, we used low-coverage WGS data from 26 animals caught in Brazil (*26*) The final WGS-derived genotyping data for the marmosets used in most downstream analyses consisted of roughly 60 million single nucleotide variants from 810 individuals (excludes the low-coverage animals from Brazil).

### Cryptic population structure in marmosets

Population structure, or a pattern of systematic differences in allele frequencies between subpopulations, arises due to non-random mating. From a practical standpoint, delineating population structure across captive marmosets is important to avoid confounding in biomedical research. For example, there are significant differences in infectious disease susceptibility, disease pathogenesis and behavior between Indian-origin and Chinese-origin rhesus macaques, despite their moderate differentiation (FST∼0.15) (*43*, *44*).

Beyond experimental concerns, population structure can also reveal the evolutionary history of marmosets in the wild. The common marmoset is native to the Northeast region of Brazil, where it occupies remarkedly distinct biomes, ranging from the humid Atlantic Forest to the semi-arid Caatinga (*24*). However, given the scarcity of genetic data for this species, little is known about population structure and differentiation in the wild. Indeed, there are not even robust estimates of the census population size of wild marmosets.

First, we investigated population structure by performing dimensionality reduction on the full genomic dataset of 805 captive marmosets (**Fig. 1D, Figs. S2-S3)**. We found that, in PC space, the largest axis of variation represents the differences between animals born at CIEA & CLEA (both colonies in Japan) and NEPRC (the now extinct New England Primate Research Center). Utah also forms a distinct cluster, likely reflecting limited gene flow from other US colonies. Most other US colonies overlap in PC space, indicating that animals have been exchanged, or that the colonies were derived from genetically similar ancestral sources.

PCA also captures recent strong familial structure, for instance caused by strong inbreeding within a particular colony. Thus, it is not clear to what extent the PCA reflects recent, post-colony formation history or ancestral population structure in natural populations. To disentangle these, we computed FST, a measure of relative differentiation, for all pairs of colonies using unrelated individuals (**Fig. 2A**). We found that most colonies are only weakly differentiated, with FST values ranging from 0.0 to 0.05. Surprisingly, the largest differentiation between CIEA and NEPRC reaches an FST of roughly 0.15. This amount of population structure is comparable to what is seen between humans from Europe and Africa (*45*).

**Figure 2.**
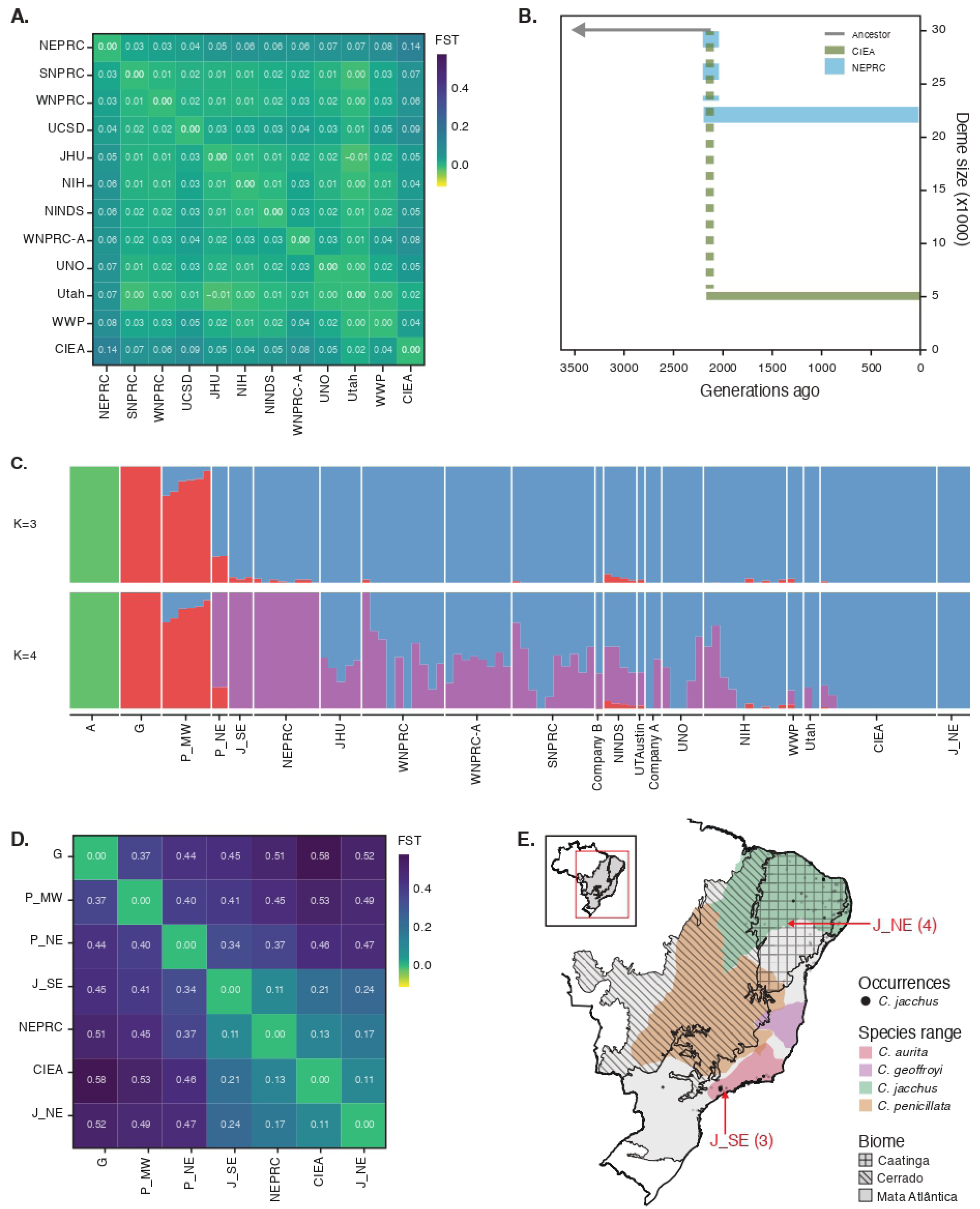
Population substructure across captive marmosets and relationships with marmosets sampled in Brazil. (A) Pairwise FST between captive colonies. Only unrelated individuals were used. (B) 2-population split model that best fits the joint site frequency spectrum of NEPRC and CIEA. (C) Admixture proportions estimated using NGSAdmix on unrelated MCC samples (down sampled to 4x) and samples collected in Brazil (all low coverage). A = *C. aurita*, G = *C. geo,royi*, P_MW = *C. penicillata* from the Midwest of Brazil, P_NE = *C. penicillata* from the Northeast, J_SE = *C. jacchus* from the Southeast and J_NE = *C. jacchus* from the Northeast. (D) Pairwise FST between di<erent *Callithrix* species and the two MCC colonies on the opposite ends of the axis of largest variation, NEPRC and CIEA. (E) Map with ancestral ranges of *Callithrix* species, the biomes the ranges overlap with, and occurrences data from GBIF (black dots). The sampling locations of *C. jacchus* obtained in Brazil are highlighted with red arrows.

To better understand the differentiation between CIEA and NEPRC, we fitted a simple 2-population split model to the observed joint site frequency spectrum using dadi-cli (*46*, *47*). Importantly, this accounts for post-split population size reductions (*e.g.*, driven by a founder effect in captive populations or recent inbreeding), which could also increase differentiation. Under idealized conditions, we estimated that these two populations split from a recent common ancestor around 2,100 generations or 10,100 years ago (**Fig. 2B**). This split predates the formation of captive colonies by millennia, providing clear evidence for population structure that must reflect differences across wild marmoset populations.

We next sought to characterize the captive marmoset genomes in the context of animals sampled in Brazil. We found that NEPRC is more closely related to animals from the southeast of Brazil, whereas CIEA is more genetically like individuals sampled in the species’ native range (northeast) (**Figs. 2C-D**). This supports the hypothesis that the NEPRC and CIEA were founded from genetically different wild sources in Brazil, perhaps reflecting the different biomes, the Atlantic Forest and Caatinga, where the common marmoset can be found (**Fig. 2E, Fig. S4**). We also found that some captive animals contain low-levels of *C. penicillata*-like ancestry, with NINDS showing the highest admixture proportion (∼4%, **Fig. 1C**). However, without a comprehensive understanding of genomic variation within *C. jacchus* and across other *Callithrix* species, it is hard to pinpoint the exact location of these different population sources.

### Genetic variation within colonies reflects ancestry and informs colony management

Together with genetic structure, levels of genetic diversity within colonies give clues into evolutionary history and inform breeding and colony management. First, we computed heterozygosity, the number of heterozygous sites per callable base pair, for each individual prior to imputation (**Fig. 3A, Fig. S6**). We observed a median of 1.28 heterozygous sites per kilobase (kb), with an inter-quartile range of 1.19 to 1.4. This heterozygosity is comparable to humans in Africa, but notably lower than in rhesus macaques, which generally ranges from 2 to 3 heterozygous sites per kb (*48*). There was considerable variation in median levels of heterozygosity across colonies, with NINDS and NEPRC harboring the most variation (median of 1.51) and CIEA the least, with a median of 0.91. Heterozygosity was correlated with a putatively ancestral source population from the southeast of Brazil, which was more closely related to NEPRC (**Fig. 3F**). Beyond having high heterozygosity, NEPRC showed substantial variation in levels of genetic diversity along the genome (**Fig. S8**).

**Figure 3.**
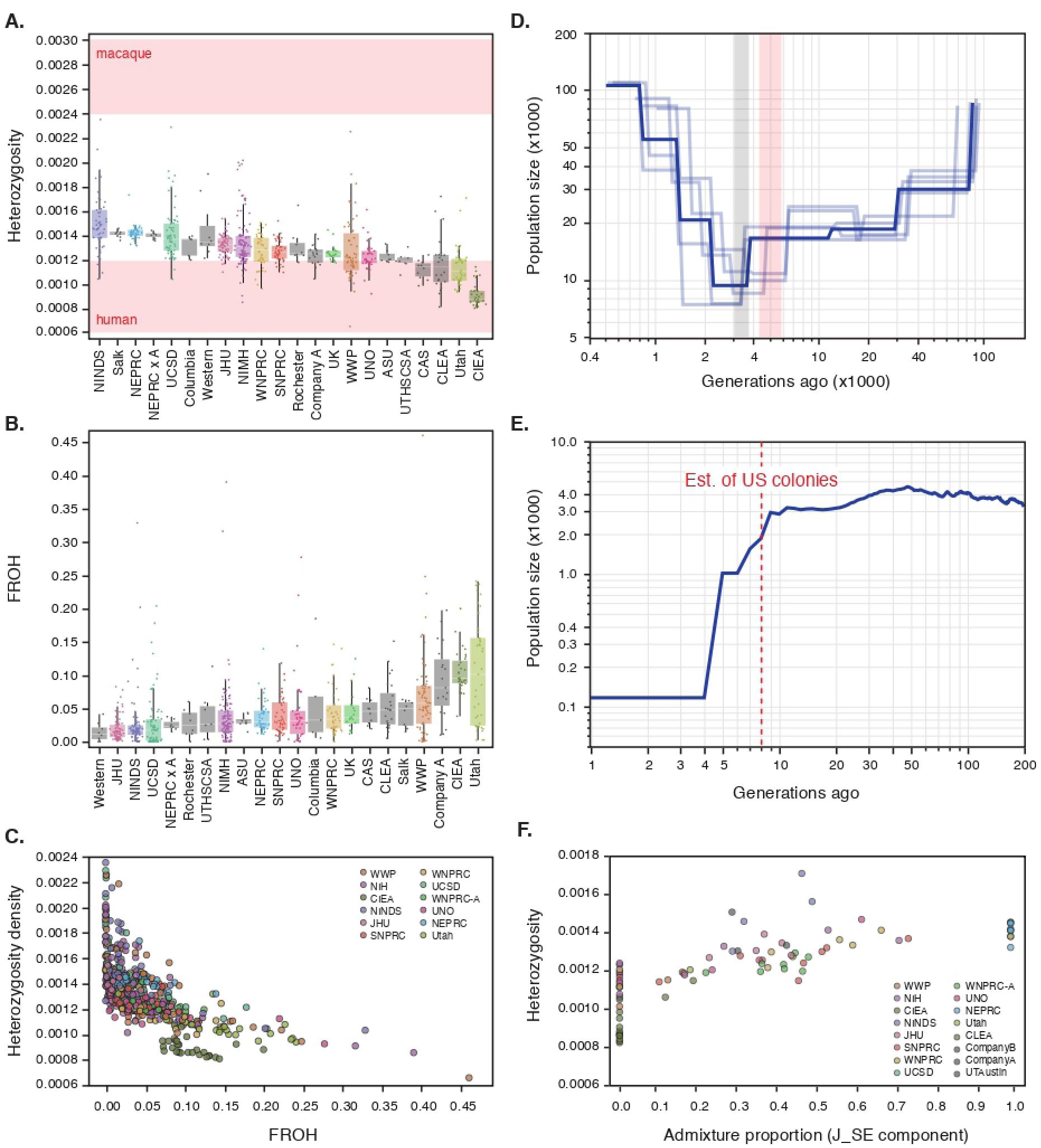
**Genetic variation across marmoset colonies and demographic history**. (A) Heterozygosity across marmoset colonies (excluding variants with minor allele count less than 2); shaded red areas show range of heterozygosity seen in humans and rhesus macaques. (B) Fraction of the genome covered by long runs of homozygosity (FROH, >1Mb) across individuals. (C) Relationship between heterozygosity and FROH. (D) Inferred long-term demographic history of marmosets using SMC++. Only a subset of unrelated individuals that formed a homogeneous cluster were used, to mitigate the impact of recent familial structure as well as population substructure. The Last Glacial Maximum (LGM) is shaded in red, and the Heinrich Stadial 1 (HS1) is shaded in light gray. (E) Inferred recent demographic history of marmosets using GONE. The dashed red line indicates the approximate time when marmoset colonies started in the US. Same set of individuals as used in C. (F) Relationship between heterozygosity and the proportion of J_SE component (in green; from Figure 2C).

To examine the amount of inbreeding across colonies, we identified runs of homozygosity (ROH) in each sample. Large stretches (or runs) of homozygosity arise when closely related individuals breed, thus inbreeding can be estimated by computing the fraction of the genome covered by such runs of homozygosity, or FROH (*49*, *50*). We observed a median FROH of 2.85%, with an inter-quartile range of 1.45% to 5.92% (**Fig. 3B**). About a third of the samples showed at least moderate levels of inbreeding, with FROH surpassing 5%, and roughly one in twenty samples showed signs of strong inbreeding, with FROH greater than 15%. There was significant variation in inbreeding across colonies, with CIEA and Utah individuals having the highest inbreeding coefficients, with a median of 9.59%.

A high FROH can be caused either by recent inbreeding in captivity or low effective population sizes in wild marmoset populations. By looking at the length of ROHs, it is possible to estimate the number of generations since a shared ancestor (*51*). Under a simplified model and assuming a recombination rate of 10^−8^, a run of length greater than 5Mbp implies a shared ancestor roughly 10 generations ago. We found that Utah individuals contained the greatest number of long (>5Mb) ROHs, with a median of 14.5 long ROHs per individual (**Fig. S9**). Only for Utah can the inbreeding signals be explained by their captive history (in the last 10 generations or 48 years).

Although CIEA also has elevated levels of FROH, most of the ROHs are of intermediate length (∼1Mb) and had an ancestor about 50 generations (or 240 years) ago. Indeed, CIEA animals have disproportionately low heterozygosity given the observed levels of inbreeding (**Fig. 3C**). Taken together, these results support the conclusion that there must exist strong population structure in wild marmoset populations that predates the establishment of research colonies, and that CIEA founders come from a different but smaller natural population.

Although molecular estimates suggest ancestral inbreeding may be moderate to high across portions of the captive marmoset population, available pedigree data does not indicate wide-spread recent consanguineous mating. Rather, the inbreeding coefficient is bound to increase due to smaller population sizes in captivity. Targeted breeding practices can effectively limit future inbreeding: for example, we saw a reduction in inbreeding when individuals from two source colonies, NEPRC and Company A, were crossed. The two source colonies had a median FROH of 3.13% and 8.10%, respectively, whereas their offspring exhibited a lower inbreeding coefficient of just 2.29% (**Fig. 3B**). This demonstrates that cross-colony breeding can effectively reduce genomic inbreeding and supports the further development of cooperative breeding management across captive colonies.

### Evidence for recent and ancient population bottlenecks

The population genomic data can also be used to clarify the recent and ancient demographic history of captive marmosets. We used GONe to estimate very recent effective population size changes, a method that relies on linkage disequilibrium (LD) patterns (*52*). We found a sharp decline in size that coincides with the establishment of colonies in the US, roughly 50 years ago. Currently, the effective population size of captive marmosets is approximately 100 individuals (**Fig. 3E**). Tracing lineages back in the pedigree records yields approximately 400 founders, though this number is probably an overestimate given the patchiness of the historical records.

With such small effective population sizes, the captive population is bound to lose genetic diversity quickly due to genetic drift and inbreeding (*53*). Thus, it is essential that colonies coordinate breeding strategies to minimize the effects of inbreeding going forward.

We also investigated more ancient demographic events by inferring an effective population size trajectory using SMC++, a method that relies both on the site frequency spectrum and LD information of traditional coalescent hidden Markov models (*54*). We estimated that the common marmoset experienced a bottleneck that cut its effective population size in half, from 20,000 to 9,000 individuals, nearly 20,000 years ago (Figure 3D). This bottleneck coincides with two major climatic phenomena that affected South America: The Last Glacial Maximum (LGM), a period marked by significantly cooler and drier conditions in Brazil (*55*), and Heinrich Stadial 1, which followed the LGM and brought continued cooling (*56*, *57*). Such dramatic climate shifts likely contributed to habitat contraction for the common marmoset, consistent with broader patterns of habitat loss and megafaunal extinctions in South America and other regions (*58*).

More recently, we infer a steep population expansion, with the current effective population size reaching ∼50,000 individuals. It is hard to interpret this recent increase without more comprehensive knowledge of Callithrix genomic variation in the wild, though it is possible this either reflects population subdivision and subsequent admixture in the wild, perhaps between the two ancestral populations we uncovered above. Alternatively, this could be driven by recent admixture between *Callithrix* species, due to the introduction of *C. jacchus* into the range of *C. penicillata*, *C. geoffroyi* and *C. aurita* in southeast Brazil (*27*, *28*).

### Deleterious mutation burden

Deleterious mutations can reduce the fitness of individuals and affect the health and viability of a population, a cumulative effect over the population known as genetic load (*59*). Genetic load can be produced by many evolutionary processes, such as genetic drift, admixture, and inbreeding. In particular, inbreeding increases homozygosity and can expose recessive deleterious alleles, thereby elevating the genetic load (*49*). Given the pervasive signs of genomic inbreeding identified in the captive marmoset population, it is essential to quantify and compare genetic loads across different colonies.

We annotated the segregating variants as putatively deleterious using SIFT, a tool that predicts whether an amino acid change affects protein function (*60*). For each individual in our reference panel, we computed the proportion of derived deleterious alleles (out of all polarized mutations) they carry as a proxy for genetic load (*61*, *62*). We found some variation in genetic load across colonies, with NINDS animals carrying the least amount of deleterious variation and CIEA individuals the most (Figure S10). Utah, the colony with the most recent inbreeding (**Fig. S9**), presented intermediate levels of genetic load. Thus, it seems that most of the differences in load stem from the different wild origins of the CIEA animals, and recent inbreeding has not affected load. This is in accordance with what has been seen across human populations (*63*), as genetic load may not necessarily increase with a reduction in effective population size if consanguineous mating is avoided.

### Genetic maps indicate reduced recombination in marmosets

Recombination rates can vary significantly along chromosomes, depending on distance to telomere, GC content, and recombination hotspots (*64*, *65*). Understanding this variation is crucial for establishing the marmoset model, because estimates of recombination rates are a key parameter in genetic and evolutionary analyses (*66*, *67*). Here, we estimated a fine-scale recombination map for marmosets using both pedigree and linkage disequilibrium information (See Supplemental Material).

We identified 740 crossover events from 36 meiosis, yielding a sex-averaged genetic map length of 2,055 cM. The marmoset genetic map length is about 15% shorter than that of macaques and 40% shorter than the human map (*68*, *69*). Thus, our estimate suggests a phylogenetic trend towards reduced genetic map in primates more distantly related to humans (whose map is about 3500 cM). In accordance with this trend, lemuriform primates (strepsirrhines) possess an even shorter genetic map (*70*).

We observed a sex-averaged recombination rate of 0.79 cM/Mb, considerably lower than the rate for humans, which is 1.13 cM/Mb (*69*). One possible factor contributing to a lower recombination rate is increased genome stability. The marmoset genome accumulates *Alu* elements at a slower pace than humans and macaques, pointing to greater stability in marmosets (*34*, *71*). A reduced recombination rate may have evolutionary consequences. For example, the lower recombination rate in marmosets may explain the observed lower levels of genetic diversity, because the linked effects of natural selection are expected to be stronger (*72*).

The ratio of female to male genetic map length was close to 1.1 (2150 cM vs. 1961 cM), which is lower than that of humans (1.36) (*69*), but more like the ratio observed in rhesus macaque (1.2) and a lemuriform primate (1.15) (*68*, *70*). The shorter generation times for the non-human primate species may explain the smaller differences between male and female map lengths, as there is less opportunity for double strand breaks to occur during oocyte arrest (*73*).

We also estimated a fine-scale population recombination map by analyzing linkage disequilibrium patterns across unrelated individuals using pyrho (**Fig. 4A**) (*74*). We estimated an average population recombination rate of 1.5 × 10^!#^ per bp per generation (or 0.15 cM/Mb). It is not trivial to convert the population recombination rate (4𝑁_$_𝑟) into the recombination rate (𝑟 ), as this depends on an adequate estimate of the effective population size. Although pyrho considers the estimated demographic history of the sample (estimated using smc++), the very recent decline in population sizes due to captivity which have substantially increased LD cannot be properly accounted for. Thus, we linearly scaled the inferred LD-based recombination map to match the average recombination rate estimated using pedigree information. We found that population recombination rates were generally elevated surrounding the identified cross-over break points (roughly by a factor of 2) (**Fig. 4B**), like the pattern in other primates (*75*). Further, we observed an increase in recombination rates near the telomeres (**Fig. 4C**), consistent with observations for various other species (*76*). The elevated recombination rate near telomeres might explain the elevation in genetic diversity near the ends of chromosomes (**Fig. 4A**), due to a reduction in the linked effects of selection, a pattern also observed in the great apes (*67*). These observations lend support to the inferred population recombination rate map, which will prove useful for future studies using the marmoset model.

**Figure 4.**
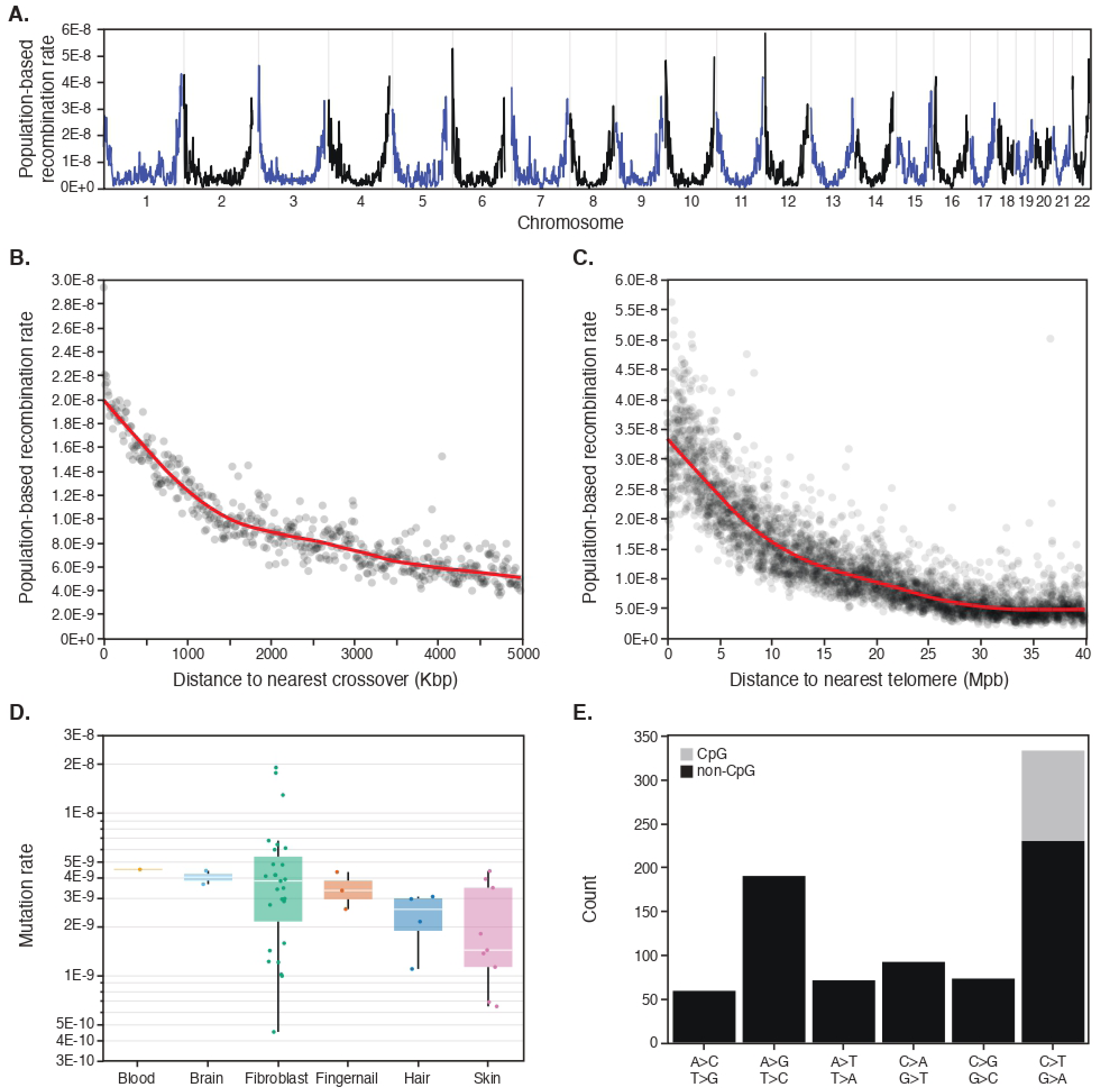
Recombination and mutation rates in the common marmoset. (A) Inferred population-recombination rates along the marmoset genome (using pyrho). (B) Population-based recombination rates surrounding cross-over events (identified using genome data for families). (C) Relationship between population-based recombination rates and distance to telomeres. (D) Inferred mutation rates for trios. Each trio is colored by the sample type of the proband (blood, brain, fibroblast, fingernail, hair and skin). (E) De-novo mutation counts for each mutation type.

### Identification of de-novo mutations

De-novo germline mutations (DNMs) are the raw material of evolution, but they also can cause diseases. It is also important to investigate mutation rate variation across primates, as it can illuminate the factors driving mutation rate evolution. Using whole-genome data for 46 trios, we sought to better characterize de-novo mutations in the common marmoset.

Although we mostly sequenced hair-follicle DNA because of the relative ease of collection and low chimerism levels, our final dataset also includes different sample types, including skin, fingernail, fibroblast culture, brain and blood. We found a two-fold difference in estimated mutation rate depending on the sample type, with hair and skin having a median rate of 1.57 × 10^!#^ per bp per generation (15 trios), and fingernail, brain and fibroblast having a median of 3.89 × 10^!#^ (30 trios) (**Fig. 4D**). As expected, we observed context dependence on the observed mutations, with the highest rates observed at CpG dinucleotides (**Fig. 4E**). A previous study estimated a mutation rate of 4.3 × 10^!#^ per bp per generation based on a single trio, which is close to our estimate using fingernail, brain, and fibroblast samples. Platyrrhine primates seem to have reduced mutation rates when compared to other, larger primates (*77–79*). Smaller primates tend to have a shorter lifespan and reach sexual maturity earlier. With fewer cell divisions in gametogenesis and less time for DNA damage to accumulate, mutation rates are expected to be generally lower in these species (*80*, *81*).

The differences in mutation rate estimates between sample types are puzzling. We speculate that sequencing depth and sample quality drive these differences. Skin and hair samples had a median callable genome length (number of base pairs with good coverage across the trio) of 1.6Gbp, whereas the fingernail, brain and fibroblast samples had a median callable genome of 2.5Gbp. It is possible there is some ascertainment bias; for example, PCR amplification is uneven across the genome, favoring GC-poor regions. However, mutation rates are generally higher in GC-poor regions (*82*). In hair and skin samples, the probability of identifying true heterozygous variants was lower than other sample types, because they had an overdispersion in allele balance distribution (i.e., true heterozygous sites were more likely to be further from the expected 50% allele balance) (**Fig. S11**). Such overdispersion is consistent with a low-level contamination in the skin and hair samples, for example from residual chimerism, DNA from other animals in the social group or the hair and skin microbiome. Although we account for false negative rates in our mutation rate estimation (Online Methods), it is possible we underestimated the false negative rates, biasing the mutation rate estimates from hair and skin samples.

Across primates, fathers pass on more de-novo mutations than mothers, possibly due to the much larger number of cell divisions throughout spermatogenesis (*83*). This leads to two patterns: (i) a male mutation rate bias and (ii) a strong paternal age effect on mutations (*77*, *84*). Using read-based phasing we found most mutations were from the mother (76 out of 119 phased DNMs, p<1×10^−3^ binomial test) implying a female bias in marmosets. Most of the phased DNMs were from fibroblast samples, which should limit chimerism as a source of artifact in our results, and any misidentified somatic mutations in the fibroblasts should be random with respect to parental chromosomes (*85*).

### A living laboratory for primate genetics

The combination of WGS and pedigree data provides an opportunity to consider the MCC colonies as a living genetics laboratory – animals can be selectively mated and measured based on genotypes of interest. To facilitate this undertaking, we performed functional variant annotation on the MCC genomes and tabulated the frequency of potential disease alleles within and across colonies (See Supplemental Material).

At a high level, gene mutability analysis shows that the number of protein-altering variants per gene is highly correlated with the total number of variants in the gene (**Fig. S12**). Many of the genes at the extreme end of calculated mutability match data from similar analyses of human genes (*86*). For example, *RYR1*, *UBR4*, *LRP1* and *CSMD1* are in the top percentile of both lists for being resistant to mutations (*RYR1* has 11 missense variants in the dataset relative to 81 synonymous mutations). On the other side of the spectrum, *AHNAK2*, *FLG*, *CMYA4* and *EYS* show tolerance to protein coding changes (46 missense variants in *EYS* compared to 68 synonymous variants).

We identified 1,498 putative loss-of-function (LoF) variants affecting 1,142 genes; of these, 164 were orthologs of genes with known Mendelian disease phenotypes in human (**Table S2**). Twenty-six of these genes are recognized as linked to developmental and/or neurodevelopmental disorders, including *SALL3*, *METTL23*, *WDR11*, *SETD5*, *SEMA6B*, *STARD9* and *NEUROG1*. Interestingly, some of these genes are known to be haploinsufficient in humans, such as *SETD5* and *SALL3*. While these LoF variants are rare in the MCC census (e.g. n=25 and n=14 carriers in *SETD5* and *SALL3* respectively), our observations raise the possibility that their severe human phenotype is not shared with marmosets, perhaps due to species differences in neurobiology. This hypothesis could be tested with careful characterization of carrier animals and/or homozygous offspring.

## Discussion

A central and unexpected finding of our work is the presence of moderate population structure across captive colonies. The differentiation between NEPRC- and CIEA-derived animals (FST ∼0.15) approaches levels observed between human continental groups and suggests that the captive population derives from at least two distinct ancestral sources. The differentiation between species in the *Callithrix* genus usually falls in the range of 0.25 to 0.7 (*26*). When placed in the context of wild Brazilian populations, these components align with geographic regions, likely reflecting long-standing structure across ecologically distinct biomes such as the Atlantic Forest and Caatinga. The reduced heterozygosity and elevated inbreeding signals in CIEA animals further support a distinct demographic history, potentially involving a historical bottleneck 240 generations ago. Together, these results indicate that the genetic structure of captive marmosets largely predates colony formation and instead reflects ancestral population subdivision in the wild.

These findings about population structure have important implications for both breeders and experimentalists. Marmoset breeding can be challenging, with high infant mortality, obesity and irritable bowel disease impacting captivity husbandry (*87–89*). Wild marmosets from different biomes may have vastly different diets, with animals in the Caatinga relying more often on tree exudates, whereas animals in the Atlantic Forest have greater access to fruits (*90*). Indeed, across *Callithrix* species there seems to be signs of dietary adaptations related to glucose metabolism (*26*). Thus, it is possible that the propensity for obesity and other health conditions is aligned with ancestry, with marmosets with Caatinga-like ancestry being metabolically adapted to lower sugar diets and therefore more susceptible to obesity on captive diets. For experimentalists, the population structure in captive marmosets could introduce additional variability to studies or decrease replicability across laboratories. Chinese and Indian-origin rhesus macaques present a similar level of population differentiation to the one we uncovered here across marmoset colonies, and these differences are known to impact immune response, for example (*29*). If comparable ancestry-associated differences exist in marmosets, it could influence behavioral, physiological and immunological traits relevant to experiments. Accounting for ancestry may thus improve research that uses the marmoset model.

The common marmoset has proven to be a powerful model for the study of behavior, neurological diseases and embryonic development. To further elevate the use of this model, we have provided here some fundamental genetic resources such as genetic maps, de novo mutation call sets, and catalogs of deleterious variants in orthologs of human disease genes. These data, combined with extensive pedigree information, establish the MCC colonies as a “living laboratory” in which naturally occurring genetic variation can be leveraged to study gene function and model disease. It is our hope that the collaboration established between US marmoset colonies, including both researchers and animal care staff, lives on as a framework for coordinated action in developing this emerging primate model.

## Data availability statement

The raw whole genome sequencing data described in this project are available from the NIH Sequence Read Archive, under accession number BioProject PRJNA1068102. Code used in the analyses presented in the paper can be found at https://github.com/mufernando/marmoset-analyses.

## Conflict of interest

The authors declare that the research was conducted in the absence of any commercial or financial relationships that could be construed as a conflict of interest.

## Acknowledgements

This work was supported by NIH BRAIN Initiative grants U24MH123696, U24MH123422, U24MH123423, and the Office of the Director of the NIH under award number P51OD011092 to the Oregon National Primate Research Center. We thank our NIH colleagues Drs. Abigail Soyombo-Shoola, Rebecca Rosen, and Christina Liu for their strategic input and support of the project. We thank Rachel Ward, Haley Hassenstab and Jeffrey French for their support in providing samples and expertise in establishing the MCC.

## List of Supplementary Materials

### Materials and Methods

Table S1: Metadata on participating centers, and animals sequenced in this study.

Table S2: Putative loss-of-function (LoF) variants identified in this study, affecting 1,142 genes. Of these genes, 164 were orthologs of genes with known Mendelian disease phenotypes in human.

### Marmoset Coordinating Center: U24MH123696

Murillo F. Rodrigues^1^, Philberta Y. Leung^1^, Jamie A. Ivy^2^, Alexandra Stendahl^1^, Karina Ray^1^, Jenna N. Castro^1^, Sam Peterson^1^, Ricardo del Rosario^3^, Simon Plösch^4^, Joanna Malukiewicz^5^, Katinka A. Vigh-Conrad^1^, Benjamin N. Bimber^1^, Jeffrey D. Wall^1^, Donald F. Conrad^1^

### Marmoset Breeding Colonies: U24MH123422, U24MH123423

Cory Miller^6^, Kaylee Cooper^6^, Xiaoqin Wang^7^, Nia Bryan^7^, Emma Easter^7^, Yang Zhang^7^, Michael Wallis^8^, Kristy Gibbs^9^, Emily Truscott^9^, Katilin Belletti^9^, Jon E. Levine^10,11^, Saverio Capuano^10^, Jenna K. Schmidt^12,13^, Corinna N. Ross^14^, Donna Layne-Colon^14^, Jeffrey Rogers^15,16^

### Marmoset Working Group

Jillian Glawe^17^, Yi Zhou^17^, Juliane Daggett-Vondras^18^, Geoffrey Stephens^17^, Marina Barto^17^, Steven Eliades^19^, Qiangge Zhang^20,21,22^, Tsephel Tenzin Thangpey^20,21,22^, Guoping Feng^20,21,22^, Krystal Allen-Worthington^23^, Aikeen Jones^23^, Heather Narver^24^, Julie Brent-Cummins^25^, Milton Herrera^25^, Rose Peterson^26^, Afonso Silva^27^, Lauren Schaeffer^27^, Gregg E. Homanics^28^, Dina-Jo Graf^29^, Jude Mitchell^29^, John Reynolds^30^, Catherine Williams^30^, Trinka Adamson^30^, Alexandra Benavente^31^, Bailey Deng^31^, Samantha N. Johnson^32^, Carmen Eggleston^32^, Lauri Nurminen^33^, Agnes Lacreuse^34^, Carrie Barr^35^, Nicholas Priebe^35^, Adam B. Salmon^36,37^, Joselyn A. Castillo^36^, Caroline Garrett^38^, Frederick Federer^38^, Alessandra Angelucci^38^, Alexander C. Huk^39,40^, Joseph Wekselblatt^41^, Allison Laudano^39,40^, Amrita Nair^42^, Anirvan Nandy^42^

^1^Oregon National Primate Center, Oregon Health & Science University, Beaverton, OR, USA, ^2^Independent consultant, Erie, CO, USA, ^3^Stanley Center for Psychiatric Research, Broad Institute of MIT and Harvard, Cambridge, MA, USA, ^4^Departement of Evolutionary Immunogenetics, University of Hamburg, Hamburg, Germany, ^5^Primate Genetics Laboratory, German Primate Center, Leibniz Institute for Primate Research, GÃ¶ttingen, Germany, ^6^Department of Psychology, University of California San Diego, San Diego, CA, USA, ^7^Department of Biomedical Engineering, Johns Hopkins University, Baltimore, MD, USA, ^8^Department of Molecular and Comparative Pathobiology, Johns Hopkins University, Baltimore, MD, USA, ^9^Animal Care & Veterinary Services, Western University, London, ON, Canada, ^10^Wisconsin National Primate Research Center, University of Wisconsin-Madison, Madison, WI, USA, ^11^Department of Neuroscience, University of Wisconsin-Madison, Madison, WI, USA, ^12^Wisconsin National Primate Research Center, University of Wisconsin-Madison, Madison, WI, 53715, USA, ^13^Department of Obstetrics and Gynecology, University of Wisconsin-Madison, Madison, WI, 53715, USA, ^14^Southwest National Primate Research Center, Texas Biomedical Research Institute, San Antonio, TX, USA, ^15^Human Genome Sequencing Center and Department of Molecular and Human Genetics, Baylor College of Medicine, Houston, TX, USA, ^16^Wisconsin National Primate Research Center, University of Wisconsin, Madison, WI, USA, ^17^Speech & Hearing Sciences, Arizona State University, Tempe, AZ, USA, ^18^Department of Animal Care and Technologies, Arizona State University, Tempe, AZ, USA, ^19^Department of Head and Neck Surgery & Communication Sciences, Duke University School of Medicine, Durham, NC, USA, ^20^Yang Tan Collective, Hock E. Tan and K. Lisa Yang Center for Autism Research at MIT, Massachusetts Institute of Technology, Cambridge, MA, 2139, USA, ^21^McGovern Institute for Brain Research, Massachusetts Institute of Technology, Cambridge, MA, 2139, USA, ^22^Department of Brain and Cognitive Sciences, Massachusetts Institute of Technology, Cambridge, MA, 2139, USA, ^23^Veterinary Medicine and Resources Branch, National Institute of Mental Health, National Institutes of Health, Bethesda, Maryland, USA, ^24^Office of Laboratory Animal Science, NIAAA DICBR, Rockville, MD, USA, ^25^Animal Health and Care Section, NINDS IRP, Bethesda, MD, USA, ^26^Bioinformatics Core, Information Technology Program, NINDS, NINDS IRP, Bethesda, MD, USA, ^27^Department of Neurobiology, University of Pittsburgh Brain Institute, University of Pittsburgh, Pittsburgh, PA, USA, ^28^Department of Anesthesiology, University of Pittsburgh, Pittsburgh, PA, USA, ^29^Department of Brain and Cognitive Sciences, University of Rochester, Rochester, NY, USA, ^30^Systems Neurobiology Laboratory, The Salk Institute for Biological Studies, La Jolla, CA, USA, ^31^Department of Biological Sciences, SUNY College of Optometry, New York, NY, USA, ^32^Committee of Computational Neuroscience, University of Chicago, Chicago, IL, USA, ^33^College of Optometry, University of Houston, Houston, TX, USA, ^34^Department of Psychological and Brain Sciences, University of Massachusetts at Amherst, Amherst, MA, USA, ^35^Department of Neuroscience, The University of Texas at Austin, Austin, TX, USA, ^36^Barshop Insititute for Longevity and Aging Studies, UT Health San Antonio, San Antonio, TX, USA, ^37^Geriatric Research Education and Clinical Center, South Texas Veterans Healthcare System, San Antonio, TX, USA, ^38^Department of Ophthalmology and Visual Science, Moran Eye Institute, University of Utah, Salt Lake City, UT, USA, ^39^Fuster Laboratory for Cognitive Neuroscience, Department of Psychiatry & Biobehavioral Sciences, UCLA, Los Angeles, CA, USA, ^40^Department of Ophthalmology, UCLA, Los Angeles, CA, USA, ^41^Department of Ophthalmology, UCLA, Los Angeles, CA, USA, ^42^Department of Psychology, Yale University, New Haven, CT, USA

## Materials and Methods

### Marmoset metadata and pedigree tracing

To create the MCC database, we requested that each participating center register their animals with the MCC by populating a table with 16 fields of key metadata on each animal. These data included demographic information like sex, weight, birthdate and birthplace, parental IDs, as well as details of reproductive and medical history. The database was validated through the identification and correction of date errors, ID errors (such as duplicate IDs), parent errors (e.g. parent was not alive at offspring birth, parent of wrong sex, etc.), and the reconciliation of other data outliers. An initial round of pedigree tracing identified gaps and inconsistencies based on the first round of metadata provided during the time of registration. We were able to obtain additional archived pedigree records from 75% of participating centers. As an attempt to further fill in gaps, we worked with individual centers to identify animals that had been shipped between colonies and map aliases between their IDs at the original and destination centers.

After genome sequencing data were available, molecular estimates of kinship, as described below, were used to further identify and resolve errors in the reconstructed pedigree. The final pedigree compiled within the MCC database was ultimately imported into the PMx software (*91*, *92*), which is used by zoos and conservation organizations to manage captive breeding programs, to generate pedigree-based genetic summary statistics for the marmoset colonies.

### Tissue acquisition and whole genome sequencing of MCC animals

All sample collections were performed under protocols approved by the Institutional Animal Care and Use Committees (IACUCs) of the participating institutions and in accordance with applicable institutional, state, and federal regulations. We developed a noninvasive protocol for the collection of hair follicles from marmosets (*41*) which were used on 578 animals from colonies across the US and Canada. Animal care staff from each colony followed this protocol to collect 3 tubes of 50 hairs per animal, which were shipped overnight on dry ice. Hair shafts were trimmed back to approximately ¼ inch in length. Hair follicle cells from each tube were lysed and pooled into a single tube per animal, and DNA was eluted from each lysate using an automated protocol on a Maxwell RSC instrument and stored at −20°C until library preparation. Sequencing libraries were constructed using the NEBNext Ultra II kit (New England Biolabs, Ipswich, MA) and sequenced on an Illumina Novaseq with a 2×150bp paired-end protocol, with a target depth of 30X per library. Total DNA mass and library complexity were used to balance the number of reads generated from each library; some libraries with low complexity were sequenced to lower mapped depth of coverage as these libraries were essentially sequenced to saturation.

### Additional Marmoset Sequencing Data

We obtained additional whole-genome sequencing data from other sources (see Table S1). An additional 193 MCC animals were sequenced in parallel, using DNA obtained from fibroblast cultures and skin punches. 108 of these were from the MIT colony (*21*), 23 from SNPRC, and 62 from WNPRC, which were sequenced at Baylor College of Medicine Human Genome Sequencing Center (*36*). 34 animals from the Chinese Academy of Sciences (CAS) and Japan (CLEA) were sequenced to high depth by Illumina WGS (*35*). These consisted of 28 fingernail samples and 6 blood samples (NCBI PRJNA807054).

For some analyses, we also included previously published data from a *C. jacchus* trio, and two other callitrichids, *C. kuhli* and *C. geoffroyi* (*93*). These genomic data were aligned to the same reference genome and processed the same way as all other samples.

### Sequence analysis and variant calling

To create a harmonized dataset, we jointly processed the raw data from all libraries, including both the published and original data, using a single pipeline. Sequence reads were aligned to the calJac4 reference genome using bwa-mem (version 0.6) (*94*). Duplicate reads were marked using Picard (version 3.2.0). SNPs were discovered using GATK (version 4.6.0.0) following the best practice recommendations: HaplotypeCaller was used on each sample individually and then unified using GenotypeGVCFs (*95*). To reduce false positives, we applied several filters to both SNPs and indels. At the site level, we removed SNPs using the following GATK VariantFiltration filter expression: ((vc.hasAttribute(’ReadPosRankSum’) && vc.getAttribute(’ReadPosRankSum’) < −8.0) || (vc.hasAttribute(’QD’) && vc.getAttribute(’QD’) < 2.0) || (vc.hasAttribute(’FS’) && vc.getAttribute(’FS’) > 60.0) || (vc.hasAttribute(’SOR’) && vc.getAttribute(’SOR’) > 3.0) || (vc.hasAttribute(’MQ’) && vc.getAttribute(’MQ’) < 40.0) || (vc.hasAttribute(’MQRankSum’) && vc.getAttribute(’MQRankSum’) < −12.5)). Further, additional SNPs were excluded based on: (i) excess depth, with DP more extreme than 98% of the sites, (ii) excess heterozygosity, above the 98% percentile, and (iii) outlier allele balance, with Phred-scaled p-values greater than 15. At the genotype level, we masked genotypes with GQ < 5.

### Sample quality control

We obtained whole-genome sequencing data for 805 individual marmosets, which varied considerably in sequencing depth (mostly driven by sample type). At lower coverage, it is difficult to accurately call genotypes. This issue can compound with the fact that some samples may suffer from residual chimerism and/or low-level contamination (e.g., from the skin and hair microbiome). Therefore, we decided to set aside samples with coverage lower than 20X to be genotyped and imputed as described below. The 234 samples with high coverage (which were not from the Asian dataset) were used as our reference panel.

We have previously shown that the hair follicle sequencing strategy is robust to chimerism (*21*, *41*), but we designed two strategies to quantify the issues caused by chimerism. First, we computed the proportion of heterozygous calls with extreme allele balance (AB) distribution (which we defined as AB<20% or AB>80%). We expect this proportion of extreme AB calls to be dependent on total sample depth, so we identified outliers as samples where the proportion of extreme AB calls is too extreme (more than 3 standard deviations away from the mean). Second, we leveraged the fact that we sequenced pairs of twins to identify residual chimerism using Census-seq (*96*).

### Phasing and imputation

Although it can be difficult to make accurate variant calls with lower depth, the existing information can be aggregated with our confident calls from the high depth samples to genotype and statistically impute variant calls. We created a reference panel with 234 moderate to high depth (>20X) samples. We phased the reference panel samples using Eagle2 (*97*). Then, we genotyped and imputed the lower coverage samples using GLIMPSE2 (*66*, *98*) We retained only the SNPs present in the reference panel and merged them with the imputed panel using bcftools merge (*99*). Three arbitrarily chosen animals were down sampled to a target depth of 4x and put through the imputation procedure to test for concordance with the original SNP call set (see Figure S13). The panel containing the imputed and reference samples will be referred to as the full panel and includes 805 individuals.

### Kinship

Understanding the kinship, or degree of relatedness, between pairs of animals is important for strategic breeding designed to limit inbreeding and retain overall genetic diversity. This knowledge is also useful in many downstream population genetic analyses that depend on accurate estimates of relatedness. We computed pair-wise kinship coefficients from both pedigree and molecular data. Pedigree kinship coefficients were computed based on the pedigree dataset curated by the MCC using the R package kinship2 (*100*).

Molecular kinship coefficients were computed using IBIS version 1.20.9 (*101*), assuming a constant recombination rate of 1cM/Mb. To increase the power to detect true segments, we kept SNPs with a minor allele frequency of at least 1% and a Hardy-Weinberg equilibrium exact test p-value (HWE) less than 0.1% using PLINK (*102*).

We generated a set of unrelated individuals using a graph-based approach. In summary, we constructed an undirected graph with all individuals and added an edge between any two individuals if their kinship coefficient (as inferred with IBIS) was greater than 0.0625. This threshold ensures that no two pairs of individuals are more than first cousins. Then, we used the maximum independent set algorithm implemented in the Python package Networkx to find the largest set of individuals which is guaranteed to not contain pairs of individuals more related than the afore mentioned kinship coefficient threshold.

### Genome accessibility

It is common to compute per base pair metrics from population genomic data. Some regions of the reference genome cannot be accurately surveyed with short-read sequencing data. There are multiple reasons why a region may be inaccessible, such as low complexity, reference genome assembly errors, and structural variation. We constructed accessibility maps for each individual (and to sets of individuals), by first identifying positions in the genome that were sufficiently covered (defined as depth greater than either 5x or 15x, depending on the downstream application), depth did not exceed 3x the mean of the individual, and where genotype quality was moderate (with a threshold of 20). For sets of individuals, this means taking the multi-way intersection of accessible regions. Further, we removed regions of poor mappability as identified by GenMap (parameters: -K 180 -E 0) (*103*). In analyses that used genomic accessibility masks, any variants that may have been identified within inaccessible regions were discarded.

### Genetic variation

We summarized genetic variation in various ways. First, we computed the heterozygosity rate for each individual, defined as the number of heterozygous single nucleotide polymorphisms (SNPs) per accessible base pair. To mitigate the effect of spurious calls, particularly on the lower coverage samples, we only considered SNPs with a minor allele count of 8 (though results were very similar without this filtering, see Figure S6). Further, we also computed two population genetic metrics across source colonies: nucleotide diversity and Tajima’s D over 100Kb non-overlapping windows across the entire genome using scikit-allel (*104*).

Heterozygosity and nucleotide diversity can be robust to recent demographic events, such as inbreeding. Thus, to better understand how endogamy and inbreeding was affecting the marmoset colonies we inferred runs of homozygosity (ROH). We used GARLIC, a model-based method with parameters: --error 0.001 --winsize 100 --resample 40 (*105*). We then estimated the fraction of the genome covered by runs of homozygosity (FROH) considering only ROHs longer than 1Mb.

### Population structure

To explore both recent and ancestral population structure in the captive marmoset population, we used principal component analysis (PCA) and the fixation index (FST). We ran the PCA using PLINK after performing LD-pruning (parameters: --indep-pairwise 100 10 0.2) and filtering SNPs with a minor allele count less than 2 and a HWE p-value less than 1e-7.

Population structure can affect population genetic analyses, but captive colonies are not natural population labels. For example, many animals currently living in the MIT colony originated from a Japanese colony (Central Institute for Experimental Animals, CIEA, which is not to be confused with CLEA, a Japanese company that supplies animals for research) or from the now extinct New England Primate Research Center (NEPRC). Thus, for most analyses we use the colony where animals were born as a population label instead of the colony where they currently reside (Figure S5).

Specifically for the demographic size history and recombination rate analyses, we sought to define the largest but homogeneous cluster of samples. We ran UMAP on the first N principal components (N was chosen such that the variance explained by them summed to 40%) to reduce the representation to two components using the package umap (*106*). Using this reduced representation of the dataset, we ran a density based spatial clustering algorithm called HDBSCAN (implemented in scikit-learn) to identify clusters (*107*). This yielded a cluster of 62 unrelated samples that were used in the SMC++ and recombination rate analyses described below.

To determine whether population structure emerged recently (i.e., with the establishment of US colonies) or in the wild, we estimated the split time between the two colonies with largest FST. We used the dadi-cli to estimate a simple split with no migration using the joint site frequency spectrum (*46*, *108*). Importantly, this allows us to model the change in effective population size since the split, which could confound the estimation of split times. The parameters to infer the demographic model were: --model no_mig --nomisid --lbounds 1e-3 1e-3 0 --ubounds 10 10 1 --optimizations 400 --check-convergence 50. Then, we identified the best fitting parameter out of 400 runs by comparing the log likelihood.

### Analysis of low coverage marmoset genomes from Brazil

To understand the relationship between captive and wild marmosets, we analyzed 26 callitrichids (including common marmosets) sampled in Brazil (*26*). These samples had considerably lower depth (average depth across samples was 2.6x) than the captive marmoset genomes, so to mitigate biases we downsampled the reads in our reference panel to 4x. Then, we ran PCAngsd on the genotype likelihoods computed using ANGSD to visualize the relationship between wild and captive marmosets (*109*, *110*). We then modelled each individual’s genome as a mixture of K putatively ancestral populations using NGSAdmix (*111*), varying the K from 3 to 4 and only including variants with minor allele frequency greater than 2.5%.

### Population size history

The genomic data from captive marmosets contain clues into the recent and ancient demographic events that shaped their history. We applied SMC++, a method to estimate effective population size history using both the site frequency spectrum and the distribution of pairwise coalescent times (*54*). We restricted the analysis to just high coverage, unrelated individuals within each of the identified population clusters (described in section <u>Population structure</u>). To minimize the computational burden, we used genotypes from the first 5 chromosomes. SMC++ uses distinguished individuals to estimate the distribution of coalescent times. Within each population cluster, we ran SMC++ using 5 distinguished individuals, defined as the samples with the highest coverage.

To estimate the recent population size history, we used GONE (*52*). GONE relies on the joint distribution of linkage disequilibrium (LD) estimates for pairs of loci over varying genetic distances to estimate recent effective population sizes. Because this analysis is computationally intensive, we randomly sub-sampled 10% of all bi-allelic SNPs in the unrelated samples in the reference panel for the first 10 chromosomes. To mitigate the effect of population structure, we only used samples from the largest identified cluster (as described in the subsection <u>Population</u> <u>structure</u>). GONE was run with default parameters.

### Recombination rates

Population genomic data can be used to estimate recombination rates across chromosomes by leveraging patterns of linkage disequilibrium. We used pyrho, a demography-aware fine-scale recombination rate estimation method that scales well with large sample sizes (*74*). We applied pyrho to a set of unrelated samples from the largest identified population cluster, totaling 62 genomes (see <u>Population structure</u>). We used the demographic history inferred with SMC++ and assumed a per generation mutation rate of 0.5e-9 (*112*).

We also inferred cross-over events leveraging high coverage sequencing data from families with at least two offspring. We follow a previously detailed strategy (*75*), but we provide a brief description here. For a target meiosis, for instance, which happened in a particular sire, we first identified informative sites: SNPs that are heterozygous in the sire and homozygous in the dam. This way, the maternally transmitted allele is always known. By comparing the alleles inherited by each of the offspring, it is possible to infer the haplotypic phase in the sire and identify potential recombination events when there is a switch in which haplotype is inherited in each of the offspring. The inferred crossover events were then manually inspected.

### Identification of de-novo mutations

Our dataset included dozens of sequenced trios, which can be used to estimate the per generation germline mutation rate. Finding de-novo mutations (DNMs) is challenging because they are rare (and rarer than sequencing errors), so careful consideration in the exact methodology is warranted (*78*). We identified DNMs using DeNovoGear as implemented within bcftools (*99*, *113*). We first obtained mpileup files for each trio using bcftools mpileup (parameters: -h 100 --indel-bias 0.75 -a AD,QS,SP). We then called variants using bcftools call (default parameters) and filtered the variants that did not pass basic quality control filters (QUAL/DP < 2 || MQ < 30 || MQBZ < −3 || RPBZ < −3 || RPBZ > 3 || FORMAT/SP > 32 || SCBZ > 3). Then, we used bcftools +trio-dnm2 to make provisional DNM calls for each trio using the DeNovoGear method (parameters: --use-DNG --dnm-tag DNM:prob). Based on best practices that are common in the field (*78*), we applied several filters to remove potentially spurious DNMs: (i) minimum depth across the trio of 15x, (ii) variants within 20bp of each other, (iii) variants in the regions of the genome that are hard to map or accurately call SNPs (see <u>Genome accessibility</u>), (iv) variants that were not heterozygous in the proband, (v) variants whose VAF was not between 35 and 65%, (vi) DNMs that were present in more than one read in either parent, (vii) variants that had a minor allele count in the entire dataset greater than 3, and (viii) DNMs with a DeNovoGear probability less than 50%.

To estimate the per generation per base pair mutation rate, we divided the number of DNMs by the number of bases that were accessible across the trio while accounting for both the false discovery rate (FDR) and the false negative rate (FNR) (*80*, *114*). For each individual, regions were deemed accessible for an individual if they had a depth greater than 15x, less than 3x the median of the individual, and had genotype quality (GQ) greater than 20. For the trio, we took the three-way intersection of all individual accessible regions. For each trio, we computed the de-novo mutation rate (𝜇_𝑖_) as

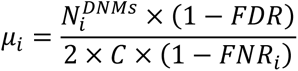

False negatives are defined as DNMs being discarded due to our strict filtering. To estimate the FNR, we first identified variants that had to be heterozygous in the offspring because both parents were homozygous for different alleles (e.g., sire 0/0 and dam 1/1). Then, we counted how many of these heterozygous calls in the offspring would be removed due to the above-mentioned VAF filter.

We manually validated DNMs by visualizing alignments of offspring, sire, and dam. We ensured that the DNM was supported by high quality reads, no evidence of structural variation in any of the individuals, and no evidence of cryptic mosaicism in either parent. Across two hundred DNMs haphazardly sampled from all trios, we could not identify any false positives, so we set the FDR to 0.5%.

Mutation rates are affected by various factors, such as mutation type, context, and parental age. First, we characterized the mutations by type (i.e., A > C, A > G, A > T, C > A, C > G, C > T). Second, we identified de-novo mutations that occurred within CpG sites. Third, we identified the most likely parent of origin for the de-novo mutations by using a read-based approach, adapted from PhaseMyDenovo (*115*).

### Deleterious mutation burden

To better understand genetic load, or the segregation of deleterious mutations, across colonies, we computed the proportion of derived deleterious alleles in each individual in the reference panel. This statistic is expected to increase in populations with a small size, where genetic drift hinders the purging of deleterious mutations (*59*, *61*, *116*).

We identified ancestral marmoset alleles using a modified version of a 447-way mammalian alignment (*42*). First, we included the reference genome calJac4 to the alignment using cactus (*117*). Then, we defined as ancestral the alleles concordant with the state inferred for the most common recent ancestor of *Callithrix* and *Callimico*. To classify alleles as deleterious or tolerated, we used SIFT, an algorithm that relies on sequence conservation (*60*).

### Functional variant annotation

Prior to functional annotation, we applied additional, strict filtering, to focus on high confidence, biologically relevant variants. We excluded 277 individuals from the master VCF: (i) outgroup species (n=5), (ii) low coverage (<10X coverage for 75% of the genome, n=242), and (iii) detectable contamination (n=30). Additionally, to filter out variants resulting from contamination (which is often sporadic) and focus on variants relevant at the population level, variants with very low observed frequency were filtered out. Specifically, the filters were: MQRankSum<-0.125, QD<10.0, AC < 5 (roughly AF ∼0.005).

We annotated the resulting VCF with SNPEff, based on MANE gene models lifted over from human to calJac4. Preliminary annotations based strictly on the calJac4 gene models showed that while most variant calls were accurate (in that they appear to represent true genetic variants), many “HIGH” impact classifications were due to predicted frameshifts or truncations of transcripts that likely do not represent the true major transcript of the gene. The affected transcripts were either minor splice variants or prediction artifacts.

## Supplemental Figures for Rodrigues Et Al

**Supplementary Figure 1.**
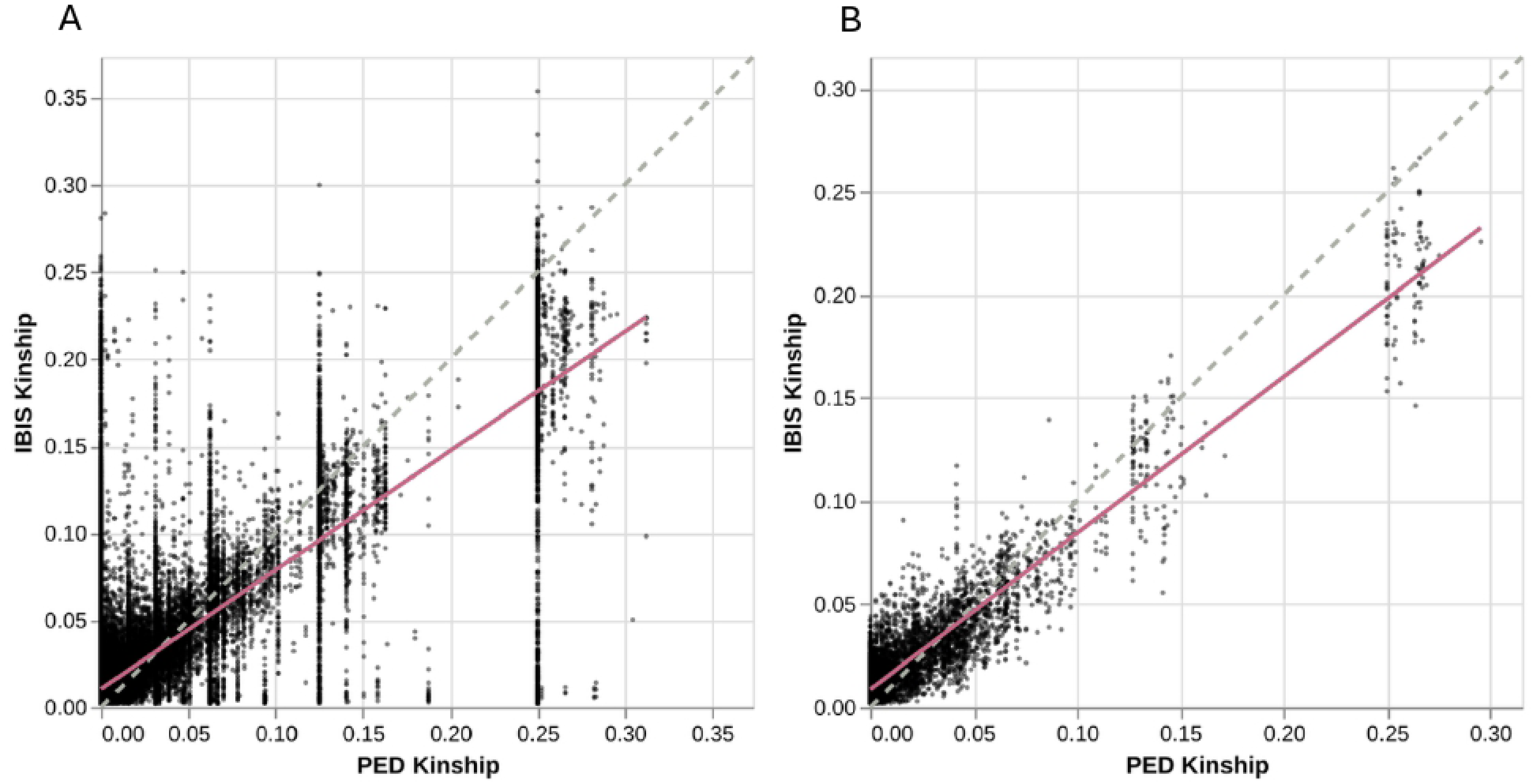
Relationship between pedigree kinship and genetic kinship. (A) Across all pairwise relationships for the 805 marmosets with genomic data. (B) Across all pairwise relationships for marmosets with pedigree depth greater than 10.

**Supplementary Figure 2.**
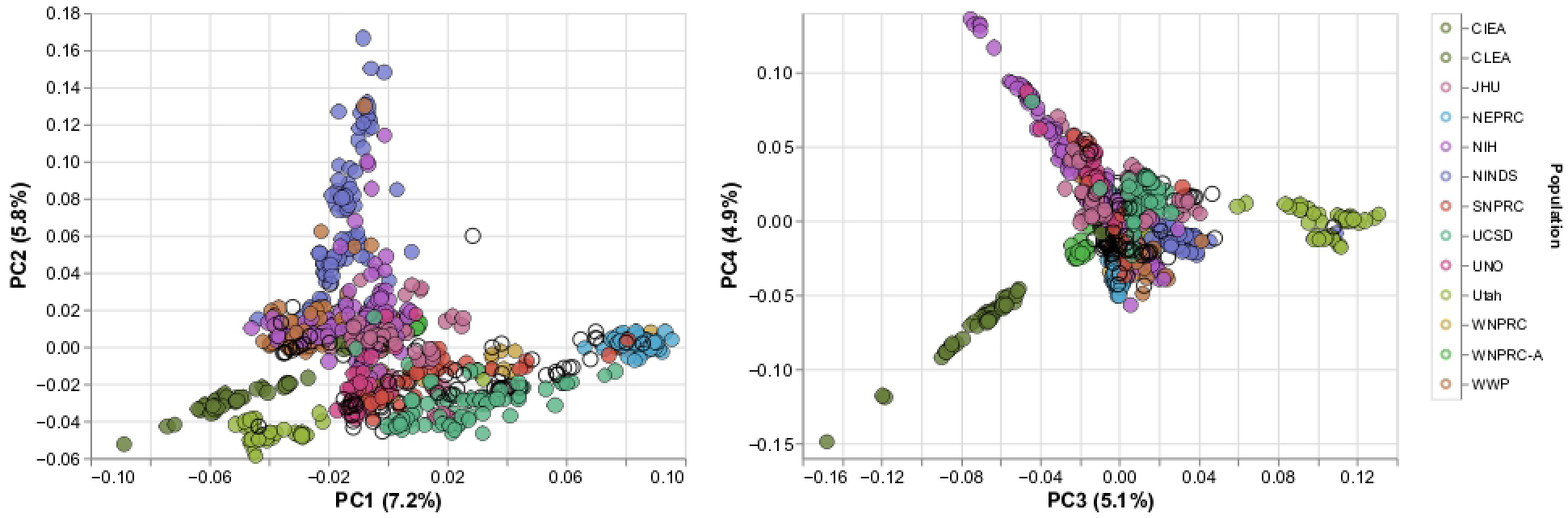
PCA for 805 marmosets using the merged panel (low coverage individuals were imputed using GLIMPSE. WNPRC-A was labelled as UK in the main paper figures.

**Supplementary Figure 3.**
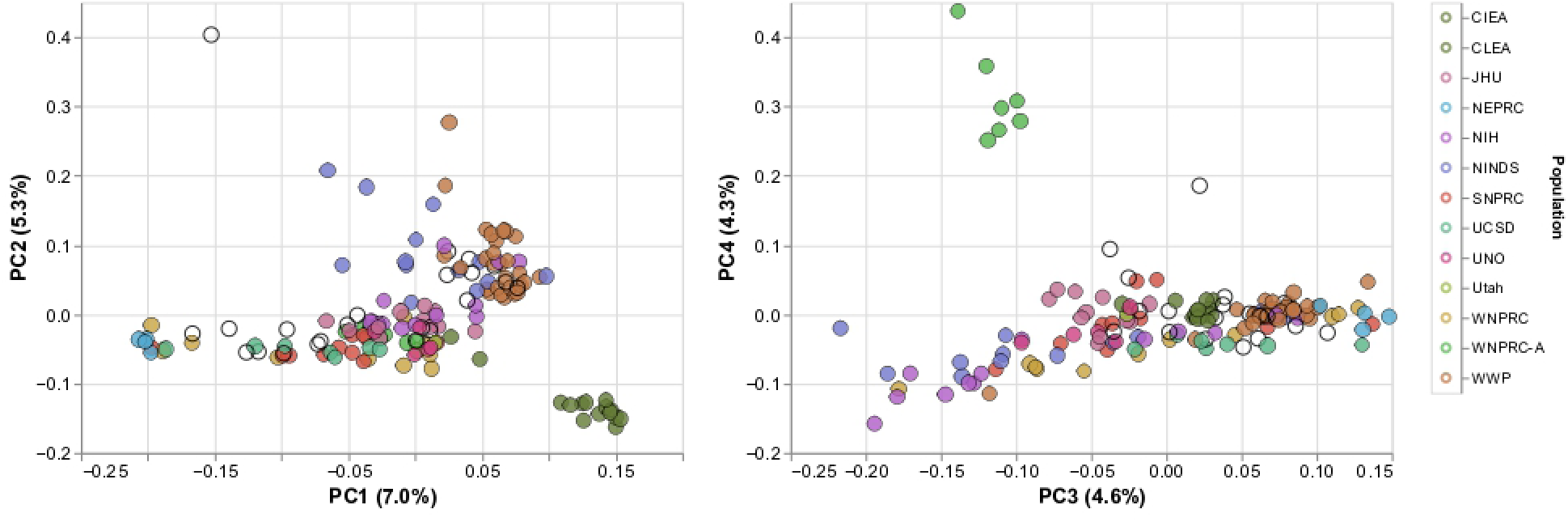
PCA for unrelated marmosets in the merged panel (low coverage individuals were imputed using GLIMPSE. WNPRC-A was labelled as UK in the main paper figures.

**Supplementary Figure 4.**
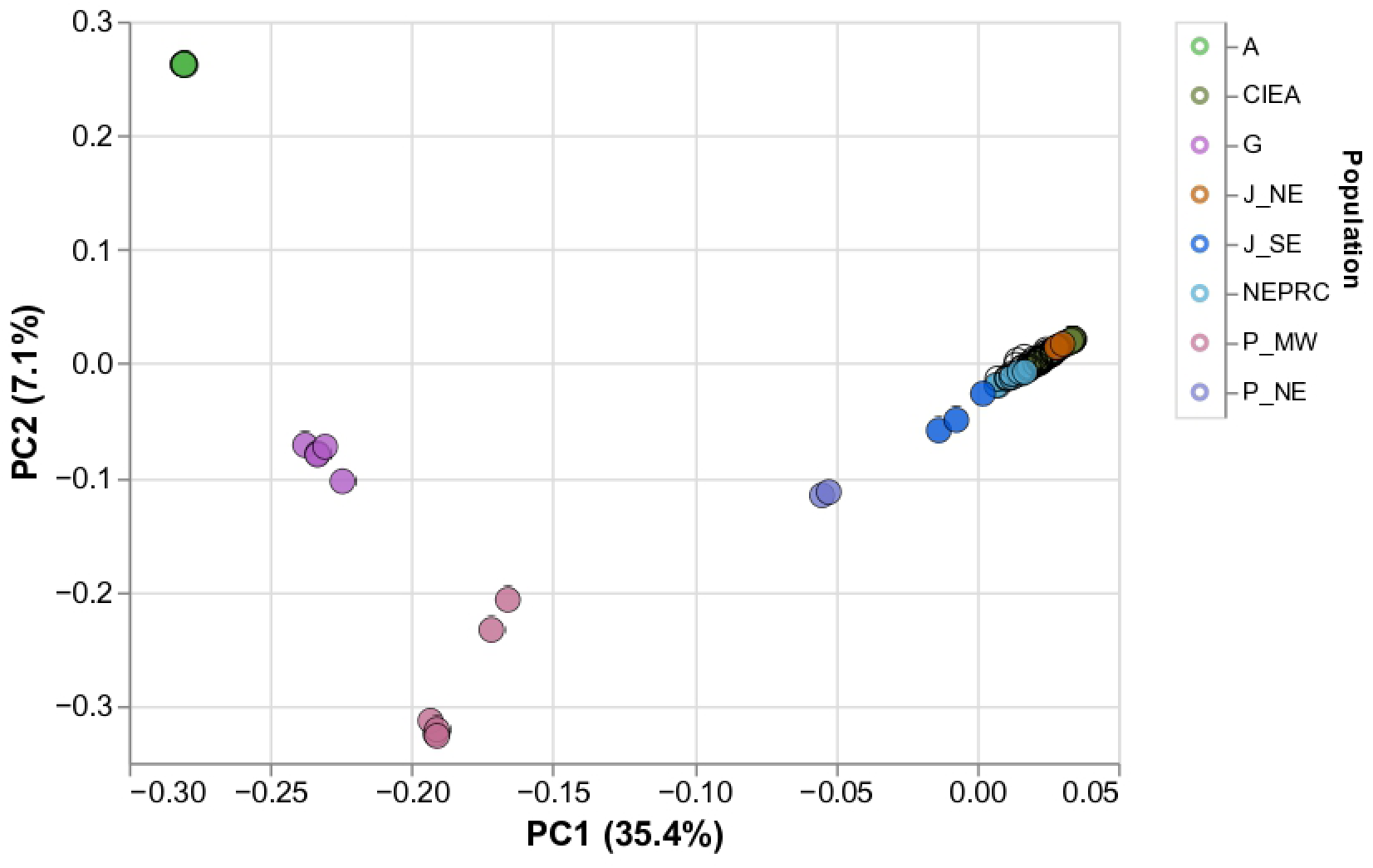
PCA unrelated MCC marmosets and 26 marmosets sampled in Brazil. All individuals were down sampled to 4x coverage (if needed), and PCA was computed using PCAngsd. A=aurita, G=geoVroyi, P_MW=penicillata from the midwest of Brazil, P_NE = penicillata from the northeast, J_SE = jacchus from the southeast and J_NE = jacchus from the northeast.

**Supplementary Figure 5.**
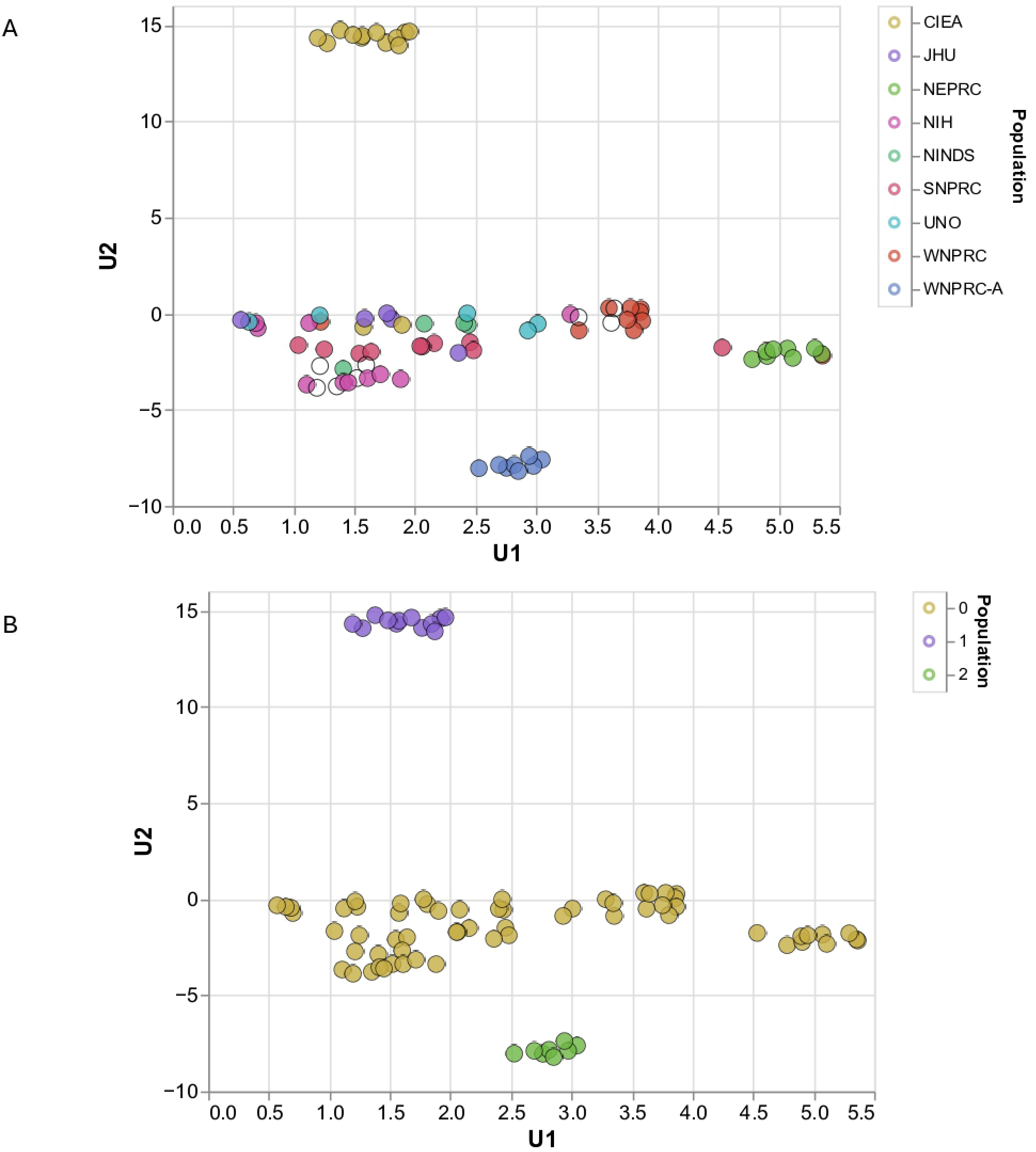
UMAP for high coverage, unrelated MCC marmosets. (A) color labels reflect source colonies. (B) color labels reflect unsupervised clusters used in downstream analyses (SMC++ and pyrho).

**Supplementary Figure 6.**
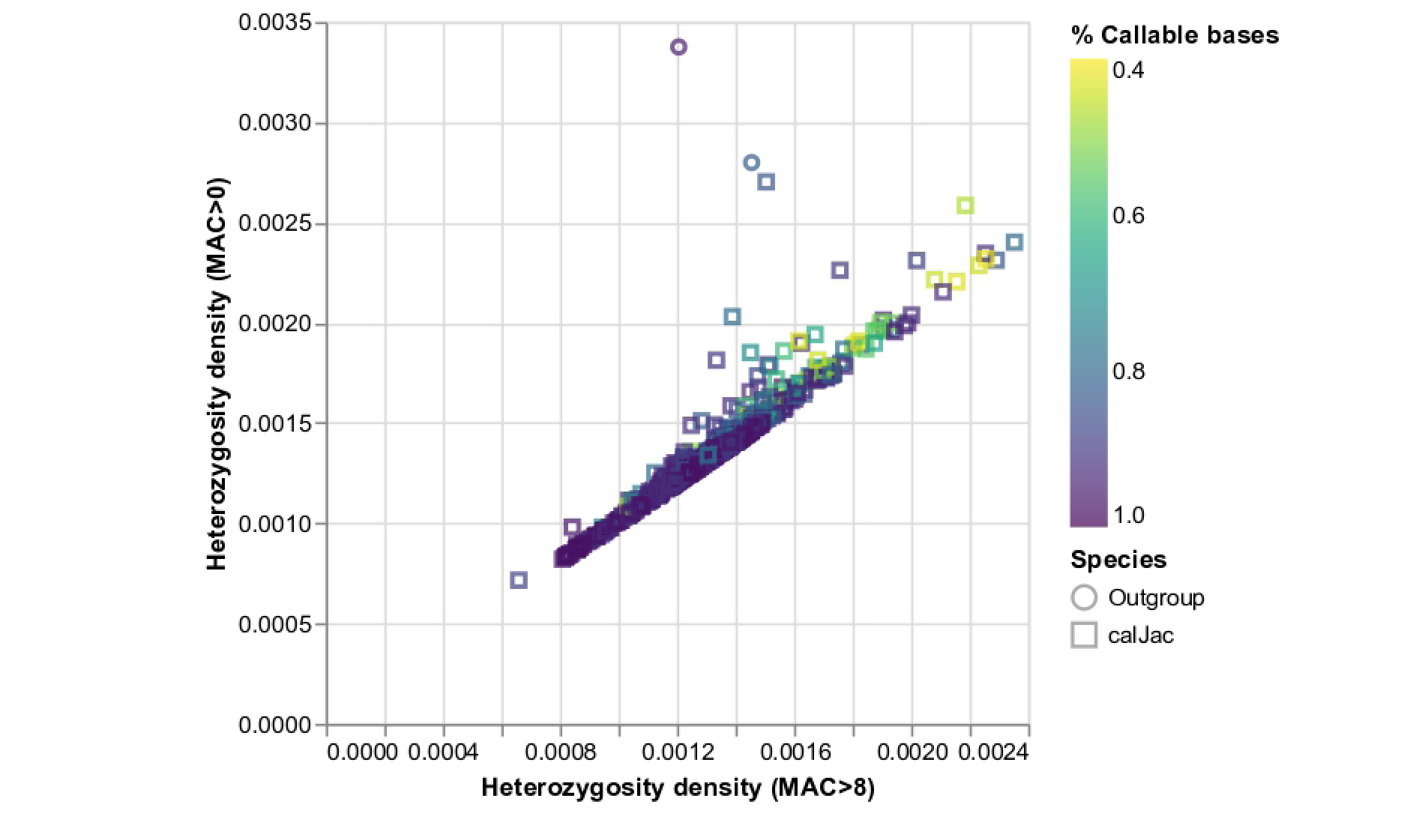
Relationship between raw heterozygosity and heterozygosity only considering variants with minor allele counts greater than 8 (roughly 1% of the samples). Two outgroup samples, C. kuhli and C. geoVroyi, are shown as circles, whereas all the C. jacchus (from MCC) samples are represented as squares. Color scale refers to the fraction of the individual’s genome has good coverage (>5X).

**Supplementary Figure 7.**
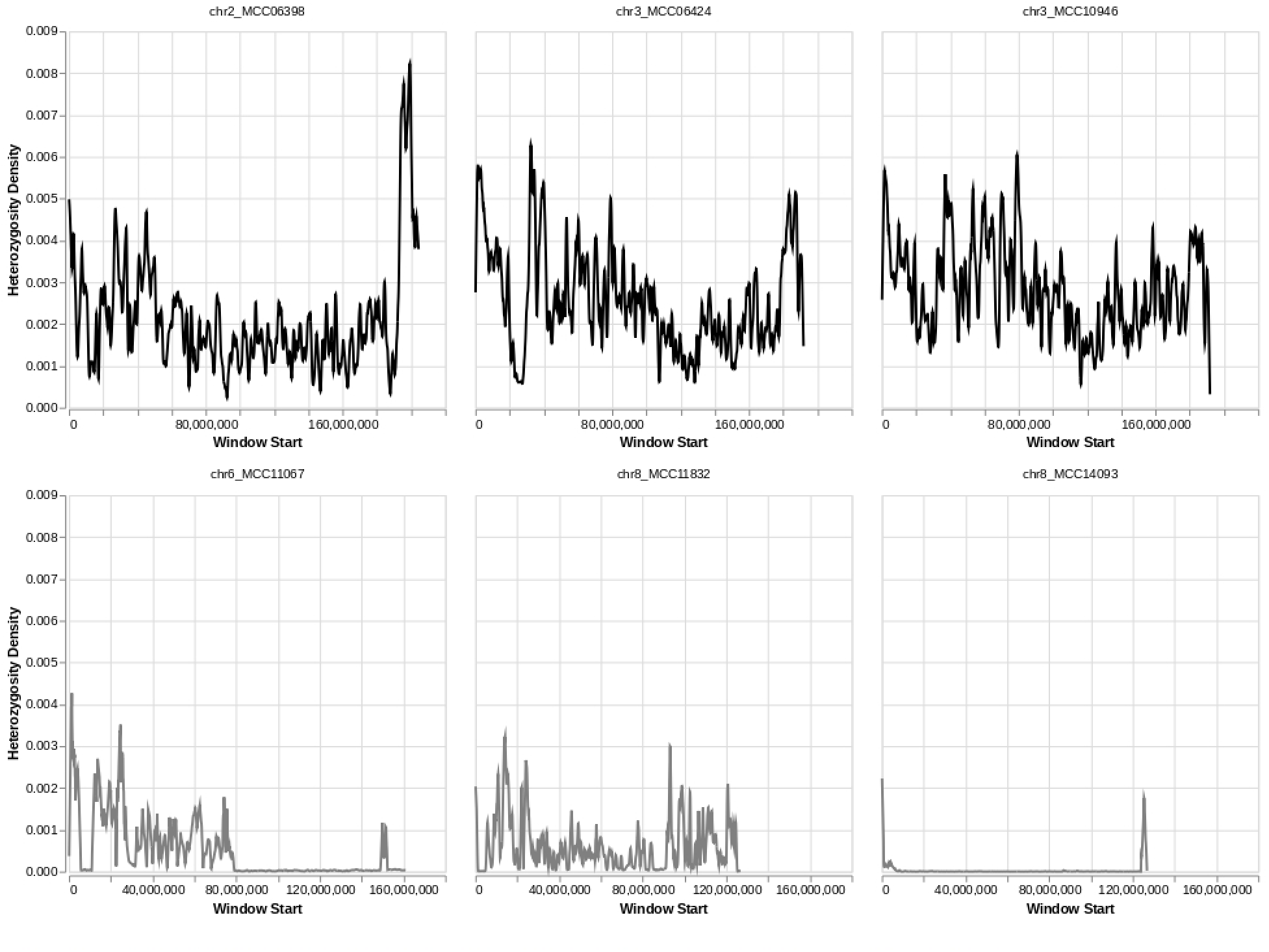
Examples of heterozygosity density along chromosomes. Top row shows individuals/chromosomes with the highest mean heterozygosity, while bottom row shows individuals with lowest mean heterozygosity. Notice regions of extreme diversity in the top row and long stretches of homozygosity in the bottom row.

**Supplementary Figure 8.**
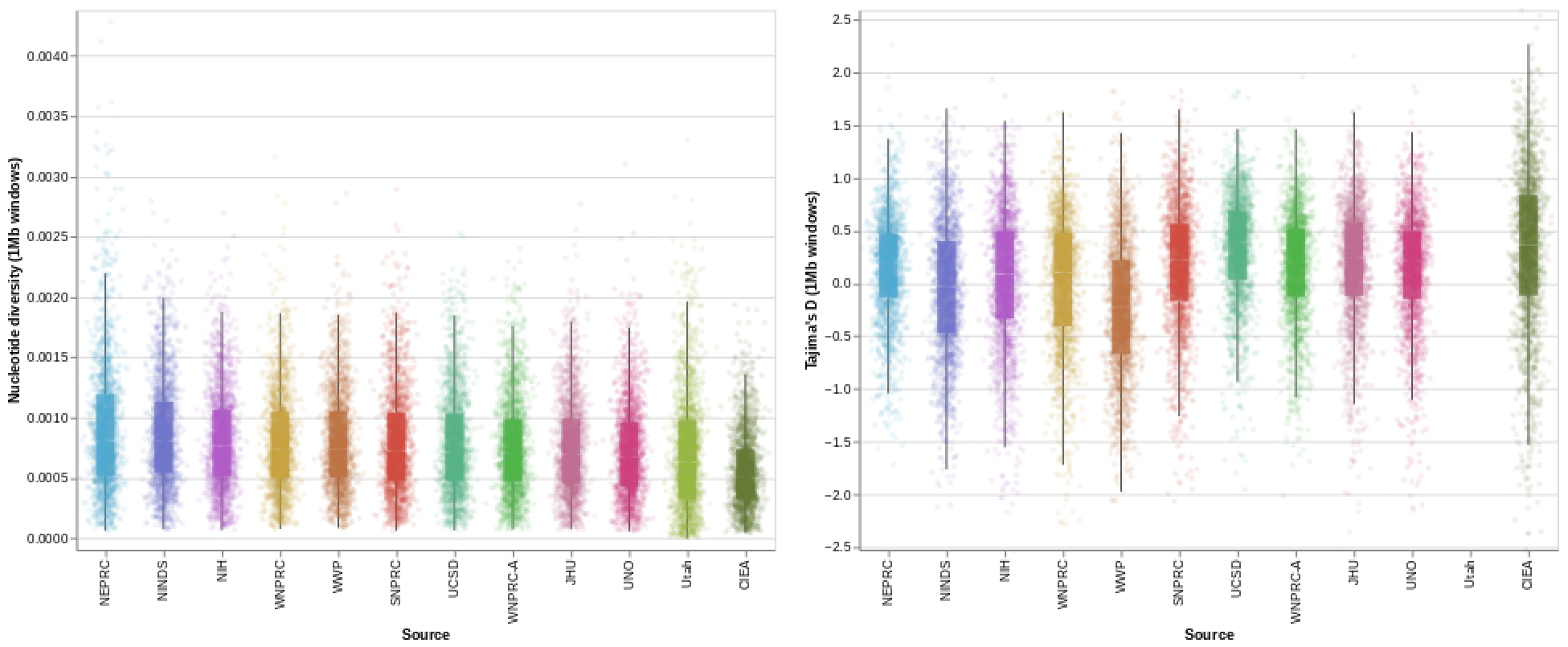
Population-genetic diversity statistics, nucleotide diversity and Tajima’s D, computed along 1Mb windows across main source colonies. Notice windows of extreme diversity (∼4x the mean) especially in NEPRC. CIEA animals show several regions with positive Tajima’s D, which is consistent with recent population contraction or bottleneck.

**Supplementary Figure 9.**
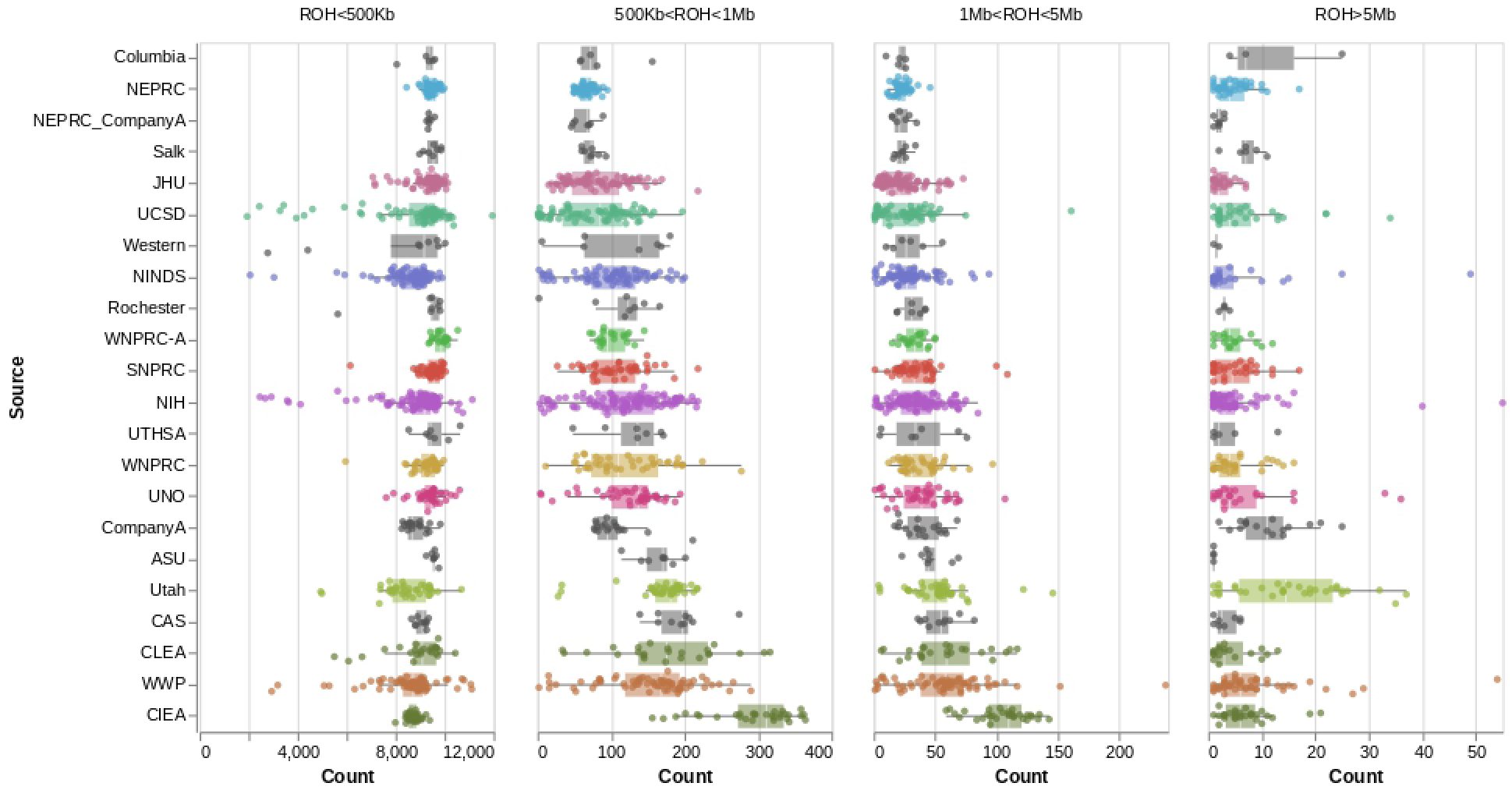
Runs of homozygosity called using GARLIC across MCC marmosets separated by run length. Notice how Utah has an excess of long ROHs (5Mb), whereas CIEA has an excess of intermediate (500Kb to 5Mb) length runs.

**Supplementary Figure 10.**
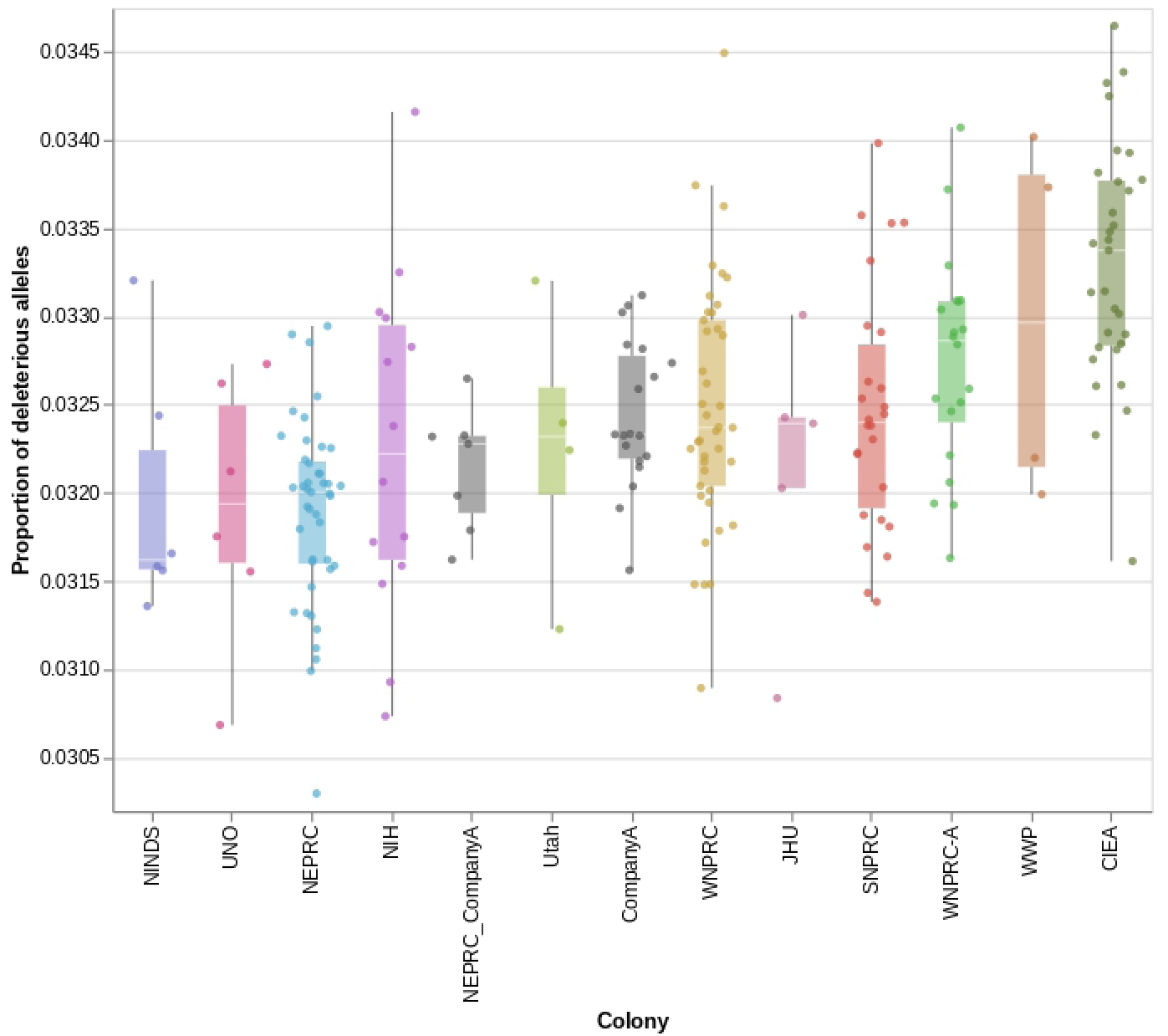
Proportion of derived deleterious alleles per marmoset (high-coverage animals only). Deleterious alleles were inferred using SIFT.

**Supplementary Figure 11.**
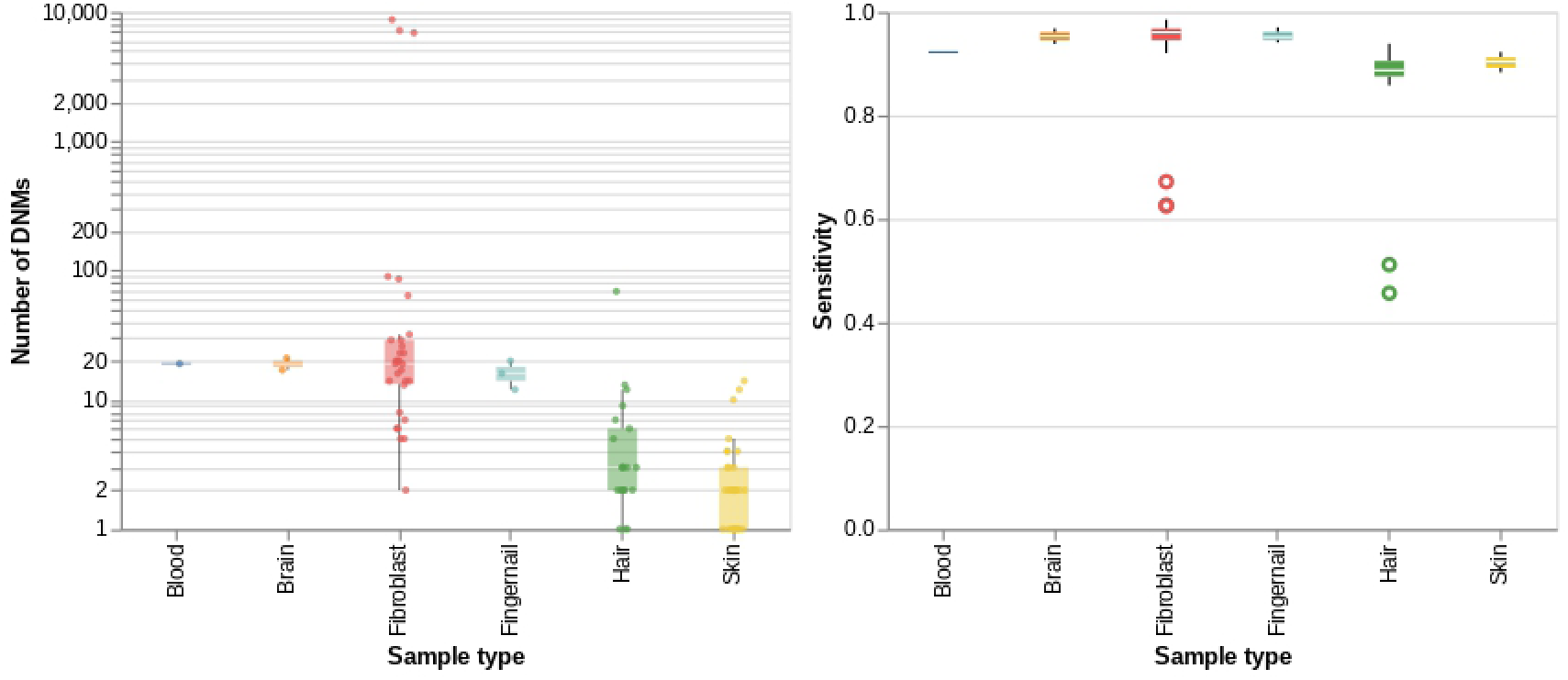
(A) Raw de-novo mutation counts across marmoset trios and sample types. Notice the extreme outliers in three fibroblast samples which likely reflect many passages during cell culture. (B) Sensitivity, or the ability of calling heterozygous genotypes when they are truly heterozygous, across sample types. Notice hair and skin had lower sensitivity, likely due to lower depth and miscalling driven by low levels of contamination. Outlier trios were omitted.

**Supplementary Figure 12.**
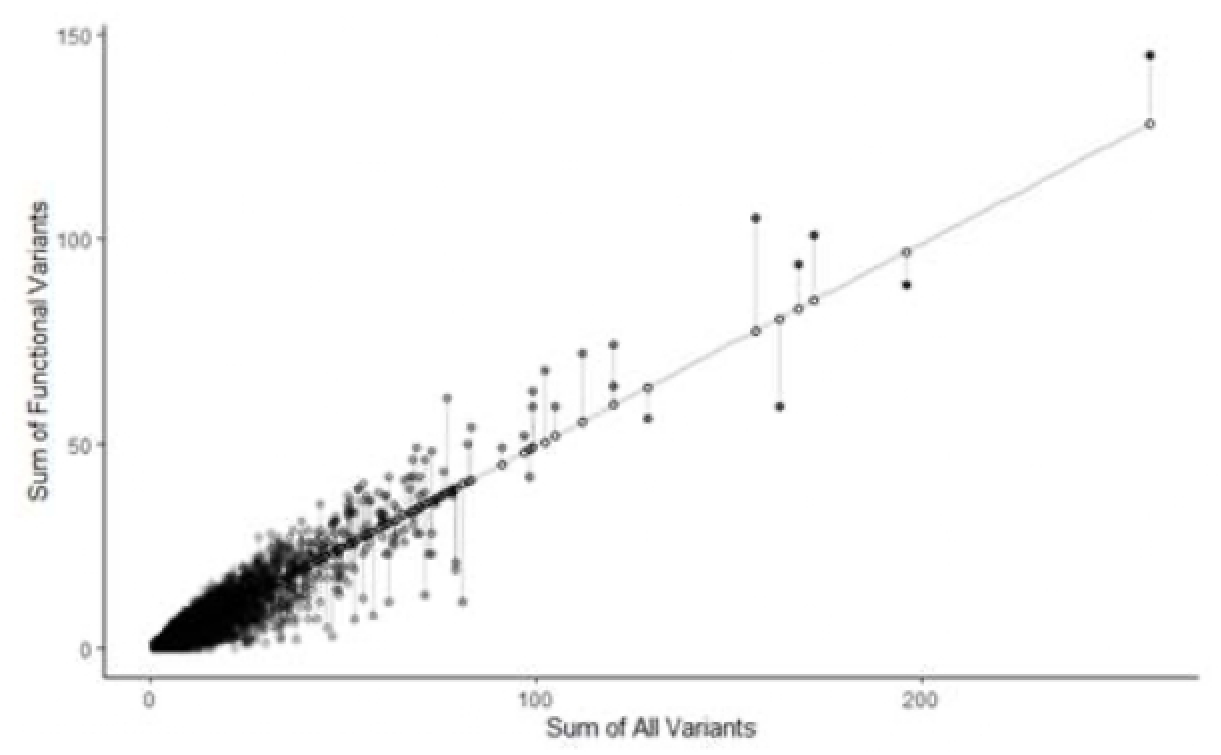
Mutability Scores Calculated with RVIS. Linear regression of the number of protein truncating and missense variants against the total number of variants observed in each gene. RVIS scores were calculated without additional filtering based on frequency. Many of the genes at the extreme end of calculated mutability match data from similar analysis on human genes. As an example, *RYR1, UBR4, LRP1* and *CSMD1* are in the top percentile of both lists for being resistant to mutations (*RYR1* has 11 missense variants in the dataset relative to 81 synonymous mutations). On the other side of the spectrum, *AHNAK2, FLG, CMYA4* and *EYS* show tolerance to protein coding changes (46 missense variants in EYS compared to 68 synonymous variants).

**Supplementary Figure 13.**
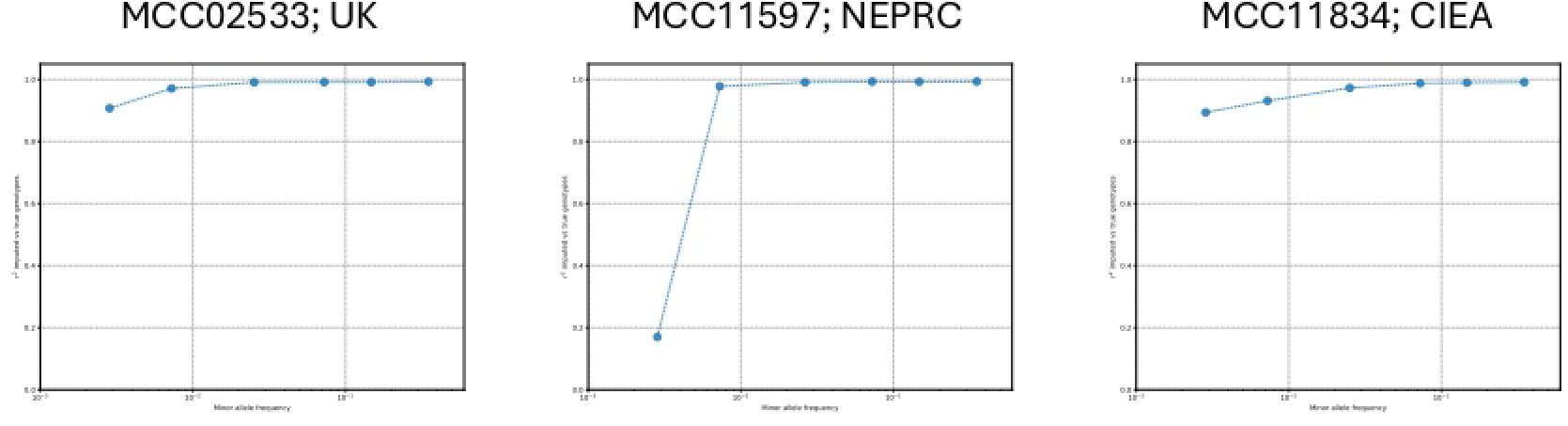
Concordance between raw and imputed genotypes in chromosome 1 across three samples at diVerent minor allele frequency thresholds.

## Notes

### Competing Interest Statement

The authors have declared no competing interest.

